# Olfactory learning modulates a neural circuit mediating innate odor-guided behavior in *Drosophila*

**DOI:** 10.1101/2023.09.20.558596

**Authors:** Florencia Campetella, Roman Huber, Martin Klappenbach, Carolin Warnecke, Fernando Locatelli, Johannes Felsenberg, Bill S. Hansson, Markus Knaden, Silke Sachse

## Abstract

Behavior is often categorized as being innate or learned, with the specific circuits being assigned to one of these categories. In *Drosophila*, neural circuits mediating an innate behavioral response are considered as being “hard-wired”, as activation of these neuronal pathways leads to stereotyped behaviors. However, only a limited number of studies assessed whether innate behaviors and their underlying neural circuits are plastic or show experience-dependent modulation. Here, we show that experience modulates second-order olfactory neurons involved in innate behavioral responses. We focus on the neural circuit defined by multiglomerular projection neurons (mPNs) that target the lateral horn, a structure relevant in the genesis of innate behavior. We show that mPNs, coding for odor attraction, are bidirectionally modulated after olfactory associative learning: when an olfactory stimulus is paired with an aversive electric shock, the activity of these neurons is decreased, while when the odor is paired with a sucrose-reward they are potentiated. We further show that this modulation requires glutamate and serotonin signaling, and that downstream third-order neurons are consequently affected. The bidirectional nature of the plasticity in these neurons is reflected in behavior: silencing mPN activity leads to odor avoidance, while artificial activation induces approach. While output from the mPNs is not required in aversive olfactory conditioning, silencing these neurons during retrieval of appetitive memories leads to a significant memory impairment. Artificially activating these neurons during odor presentation is sufficient to generate a 3 h appetitive memory. Our study in flies shows that a neural circuit coding for innate odor attraction can contribute to learned behavior, is modulated by olfactory learning and can provide reward-like reinforcement.

## Introduction

Can experience shape neuronal circuits that guide innate behavior? The vinegar fly *Drosophila melanogaster* represents an ideal system to study this question, as it displays a palette of robust innate behavior and concise underlying circuits that are traceable and amenable to manipulation. In behavioral experiments, flies show attraction or repulsion to a great number of olfactory stimuli (Knaden et al., 2012), whose detection can be sometimes attributed to a single olfactory input channel (Ebrahim et al., 2015; Kurtovic et al., 2007; Stensmyr et al., 2012; Suh et al., 2004). Artificially activating these pathways leads to a stereotyped behavior, even in the absence of an odor (Chin et al., 2018; Kurtovic et al., 2007; Suh et al., 2007). Therefore, these odor-driven behaviors are often described to be innate and the neuronal circuits that mediate them are considered as “hard-wired” (Aranha and Vasconcelos, 2018; Kim et al., 2017). However, whether experience modifies these “hard-wired” circuits has rarely been explored.

In addition to their innate behavioral repertoire, flies can also establish new associations between the elements of their environment. For instance, they can learn to associate an odor with an aversive event or reward, and modify their behavior accordingly (Quinn et al., 1974; Tempel et al., 1983). The neural circuits that are involved in the acquisition, maintenance and retrieval of olfactory memories are located in the mushroom bodies (MBs) (Boto et al., 2020; Cognigni et al., 2018). Interestingly, perturbing the MBs leads to an disruption of these memory related functions, while leaving innate olfactory behavior mostly unaffected (De Belle and Heisenberg, 1994; Heisenberg et al., 1985). Silencing neurons in the lateral horn (LH), in contrast, has been shown to alter innate behavior (Heimbeck et al., 2001; Strutz et al., 2014; Varela et al., 2019). Together, these results propelled a view in which the MBs are considered centers of olfactory learning and memory in the fly brain, with other brain areas, like the LH, governing innate odor behavior. However, previous studies showed that MB output neurons also mediate innate approach or avoidance to odors (Owald et al., 2015; Perisse et al., 2016), and that the internal state of a fly can modulate the activity of these neurons (Bräcker et al., 2013; Kadow, 2019). Conversely, recent reports showed the contribution of innate LH circuits to learned behavior (Dolan et al., 2018; Séjourné et al., 2011; Zhao et al., 2019) suggesting that the separation into innate and learned pathways (i.e. LH vs. MB) is less sharp. Therefore, we here ask whether the supposedly innate pathways undergo plasticity.

A well-characterized circuit component involved in innate behavior in the fly are the inhibitory multiglomerular projection neurons (mPNs). These neurons collect information from the antennal lobe (AL), the first olfactory processing center, and convey it mostly to the LH. Here they synapse onto third-order LH neurons (Ito et al., 1997; Jefferis et al., 2007; Lai et al., 2008; Liang et al., 2013; Okada et al., 2009; Shimizu and Stopfer, 2017). Interestingly, a subset of these mPNs respond specifically to attractive odorants and instruct innate approach behavior (Strutz et al., 2014). In addition, inhibitory input of mPNs to the LH has been shown to be selective for food odors and pheromones (Fişek and Wilson, 2014; Liang et al., 2013). We therefore asked whether this AL to LH pathway can be modulated by olfactory learning or whether this circuit is rather hardwired and exlusively encodes innate odor approach.

We used olfactory associative conditioning to show that flies can learn to avoid innately attractive odors. We determined whether learning induces plasticity in odor-evoked responses of mPNs and assessed the involvement of these neurons in the retrieval of appetitive and aversive memories. Our results reveal an unexpected plasticity in innate circuits that contribute to learnfed behavior.

## Materials and Methods

### Fly rearing

Flies were bred on cornmeal agar medium under L:D 12:12, RH= 70% and 25°C, except for experiments with TrpA1 and shibire^ts1^, where flies were bred at 23°C. For behavioral experiments, adults of both sexes were used. In calcium imaging, all experiments were conducted on mated adult females. Genotypes used in each figure are listed in the **Supplementary Table 1**. Transgenic lines were obtained from the Bloomington Stock Center (http://flystocks.bio.indiana.edu/) and Vienna RNAi Stock Center (http://www.vdrc.at). Other flies stocks were kindly provided by Kei Ito (*MZ699-GAL4*).

### Behavior

In aversive conditioning experiments, 3-5 days old flies were used and experiments were performed at 75% humidity in dim red light during training and in complete darkness during testing. Prior commencement of experiments, the odors were diluted in paraffin oil at their respective concentrations. In all aversive conditioning experiments 1% acetoin acetate (AAC), 1% benzaldehyde (BEA), 0.01% or 0.001% ethyl butyrate (ETB) dilutions in paraffin oil were used. Non-starved 3-5 days old flies were placed in a tube covered by a copper grid (CON-Elektronik, Germany). Flies received a 1 min odor presentation that was paired with 90V pulses of electic shock lasting 1.25 s at 3.75 s intervals; this was followed by a 30 s clean air interval, after which another odor (differential conditioning) or paraffin oil (absolute conditioning) was presented for 1 min in the absence of any electric shocks. Airflow was kept constant at a rate of 0.5 l min^-1^. Trained flies were immediately tested by allowing them to choose between the odors used in training for 2 min in a T-maze. The preference index was calculated based on the distribution of the flies between two compartments, normalized by the total number of flies.

In appetitive conditioning experiments, 3-5 days old flies were used. Flies were starved for 18-24 h before training and were kept starved for the entire duration of the experiment. Experiments were performed as described previously (Krashes and Waddell, 2008), and methylcyclohexanol (Sigma, 16 µl in 8 ml mineral oil) or 3-octanol (Sigma, 8 µl in 8 ml mineral oil) were used as olfactory stimuli. Briefly, groups of 50-100 flies were exposed to an odor (CS-) for 2 min followed by 30 s of room air and then 2 min of a second odor (CS+) in the presence of dry sucrose. During the test phase, flies were given 2 min to choose between two odors (CS+ vs. CS-) in a T-maze. Performance index was calculated as the number of flies in the CS+ arm minus the number of flies in the CS-arm, divided by the total number of flies (Tully and Quinn, 1985). All experiments were carried out in a climate chamber at 23°C and 55%–65% relative humidity. In experiments employing *UAS-shi^ts1^*, temperature was raised to 30°C only during the testing phase and flies were placed at this restrictive temperature 30 min before the onset of testing. In experiments involving artificial activation using *UAS-trpA1* flies were bred at 23°C and the activating temperature was set to 32°C. Temperature controls were performed at 23°C. To measure innate preference, flies were placed in the same T-maze used in conditioning experiments and allowed them to move freely between the two compartments. Innate preference was computed as the distribution of flies between the two compartments after 2 min.

### In vivo calcium imaging

Optical imaging was performed as previously described (Strutz et al., 2012). Flies were briefly anesthetized on ice and fixed with the neck onto a copper plate (Athene Grids, Plano) mounted onto a custom-made Plexiglas stage. An insect pin was used to immobilize the proboscis and the head was glued to the stage using Protemp II (3 M ESPE). Antennae were pulled forward using a fine metal wire (REDIOHM-800, HP Reid Inc). A polyethylene covered plastic slide with a small opening was placed on top of the fly’s head and sealed to the cuticle using two-component silicon (World Precision Instruments). After adding Ringer solution (NaCl: 130 mM, KCl: 5 mM, MgCl_2_: 2 mM, CaCl_2_: 2 mM, Sucrose: 36 mM, HEPES-NaOH (pH 7.3): 5 mM), the brain was exposed using a razor blade (Fine Science Tools). Cuticle, excessive fat, trachea and air sacs were carefully removed with the aid of forceps (FST). Flies trained for appetitive learning were anesthesized for <10 s 2.5h after conditioning.

After dissection flies were imaged using a Till Photonics imaging system with an upright Olympus widefield microscope (BX51WI) equipped with a 60 × Olympus objective (LUMPlanFl 60x/0.90 W Olympus). A Polychrome V provided light excitation (475nm) and a filterset ensured passage of only relevant wavelengths (excitation: SP500, dicroic: DCLP490, emission: LP515). The emitted light was captured by a CCD camera (Sensicam QE, PCOAG) with a symmetrical binning of 4(1.25 × 1.25µm / pixel). The optical plane was ∼30 µm below the most dorsal entrance point of the mPN tract into the LH (Strutz et al., 2014). All recordings lasted 10 s with a frame rate of 4 Hz. For the acquisition of the images Till Vision (Till Photonics) was used.

### Odor delivery during imaging

All odors were diluted in paraffin oil (Sigma Aldrich). Acetoin acetate (Alfa Aesar) and benzaldehyde (Acros) were used at a concentration of 1% v/v, ethyl butyrate (Sigma) was diluted to 0.1% v/v or 0.001% v/v. Odors were delivered though a custom made peek cartridge connected via Teflon tubes to a stimulus controller (Syntech). Odor pulses lasted for 2 s and had an airflow of 0.5 l min^-1^. The airflow was compensated using clean air at the same flow rate, and supplemented with 1.0 min^-1^ humidified air. A time interval of 2 min was kept between successive odor stimulations. In aversive conditioning experiments carried out under the microscope, the odor delivery was achieved using valves connected to an Analog/Digital Converter (National Instruments, USA) controlled by custom written software which was also connected to the electric shock generator (CON-Elektronik). During training, the odor stimulation lasted for 1 min, followed by a 30 s interval before the presentation of the second odor or paraffin oil for again 1 min. The odor flow was kept constant at 0.5 l min^-1^ supplemented with 1.0 l min^-1^ humidified air.

In appetitive olfactory conditioning experiments, flies were trained as described below and were dissected shortly before the imaging experiments. Dissected flies were then placed under the microscope where they were exposed to the odors for 2 s as described above.

### Imaging data analysis

Imaging data was analyzed using custom written IDL software (ITT Visual Information Solutions), kindly provided by Mathias Ditzen. All recordings were manually corrected for movement and corresponding regions of interest (ROI) were identified in each trial. In recordings from the LH, regions of interest (ROI) correspond to those ones previously identified and defined as odor-response domains in the LH (i.e. LH-PM, LH-AM, LH-AV) (Strutz et al., 2014). In recordings from vlPr neurons, the ROI was confined to processes in the LH ventral region. In AL recordings, ROIs were assigned to identified glomeruli using glomerular labeling visualized by co-expressed fluorescence of *GH-146-QF*; Q*UAS-tdTomato*. In all cases, raw fluorescent signals were converted to Δ*F*/ *F*_0_, where *F*_0_ is the averaged baseline fluorescence values 2 s before the odor onset (i.e. 0–7 frames). Odor-evoked calcium signals obtained from *MZ699-GAL4; UAS-GCaMP6s* flies, Δ*F*/*F*_0_ were calculated as the average of frames 12–20. In recordings from flies expressing *VT063948>GCaMP6s* the time window for the average was restricted to frames 10-15, as these neurons presented a faster dynamics than inhibitory mPNs. In imaging experiments from flies conditioned with a sugar reward, data from both odorants was pooled in the trained and untrained conditions. This allowed us to buffer for odor bias in individual responses.

### Immunohistochemistry

Whole-mount immunofluorescence staining was carried out as described (Vosshall et al., 2000). Initially brains were dissected in Ringer’s solution (130 mM NaCl, 5 mM KCl, 2 mM MgCl_2_, 2 mM CaCl_2_, 36 mM saccharose, 5 mM HEPES, [pH 7.3]) (Estes et al., 1996) and fixed in 4% PFA in PBS-T (PBS, 0.2–1% Triton-X). After washing with PBS-T brains were blocked with PBS-T, 5% normal goat serum (NGS). Primary antibodies were diluted in blocking solution or PBS-T and incubated at 4°C for 2–3 days. Secondary antibody incubation lasted 1–2 days.

### Statistical Analysis

Statistical analyses were performed using GraphPad Prism 7 (GraphPad). Groups that did not violate the assumption of normal distribution (Kolmogorov’s test) or homogeneity of variance (Barlett’s test) were analyzed with parametric statistics. For behavioral data, one-way ANOVA followed by Tukey’s post-hoc test was used. Significance was set at α= 0.05. One sample t tests were used to compare data against a hypothetical mean of 0. Calcium imaging data obtained in aversive conditioning experiments was analyzed with two-tailed paired t tests. Unpaired t tests were used to analyze results of flies that were functionally imaged after appetitive conditioning.

## Results

### Aversive associative learning induces plasticity in mPNs

In general, the conditioned stimulus (CS) in associative classical conditioning is meant to be neutral and only after pairing it with an unconditioned stimulus (US) is able to elicit a conditioned response (CR) similar to the unconditioned response (UR) to the US (Pavlov, 1927). The contingency of the CS and US during training assigns the valence to the CS. In *Drosophila*, odors commonly used as CS in olfactory conditioning have little or no ecological meaning to the fly. However, often odors are used in conentrations that are mildly aversive. In nature, flies are exposed to a high number of odorants to which they may be innately attracted. Can flies learn to avoid an innately attractive odorant? To answer this question, we used two associative learning paradigms: aversive absolute conditioning, which is a simple form of associative conditioning where animals learn that a single odor (CS) predicts an aversive electric shock (US) (Tully and Quinn, 1985), and aversive differential conditioning where, during training, the first conditioned stimulus is paired with an electric shock (CS+), whereas a second odor is presented alone (CS-, **Figure 1A**). As olfactory stimuli we chose ethyl butyrate, commonly found in fruits, and acetoin acetate, a byproduct of yeast fermentation, at a concentration that was equally attractive in a 2-choice assay (i.e. 1% v/v for acetoin acetate, 0.001% v/v for ethyl butyrate; **Figure 1A, B; Supplementary Figure 1A**, **B**). Flies tested immediately after the training showed a significant avoidance to the trained odor (CS+) and preferred the CS-(differential conditioning) or the solvent paraffin oil (absolute conditioning) (**Figure 1B, Supplementary Figure 1B**). We next wondered whether the neuronal circuits mediating innate attraction reflect this change in preference.

**Figure 1.**
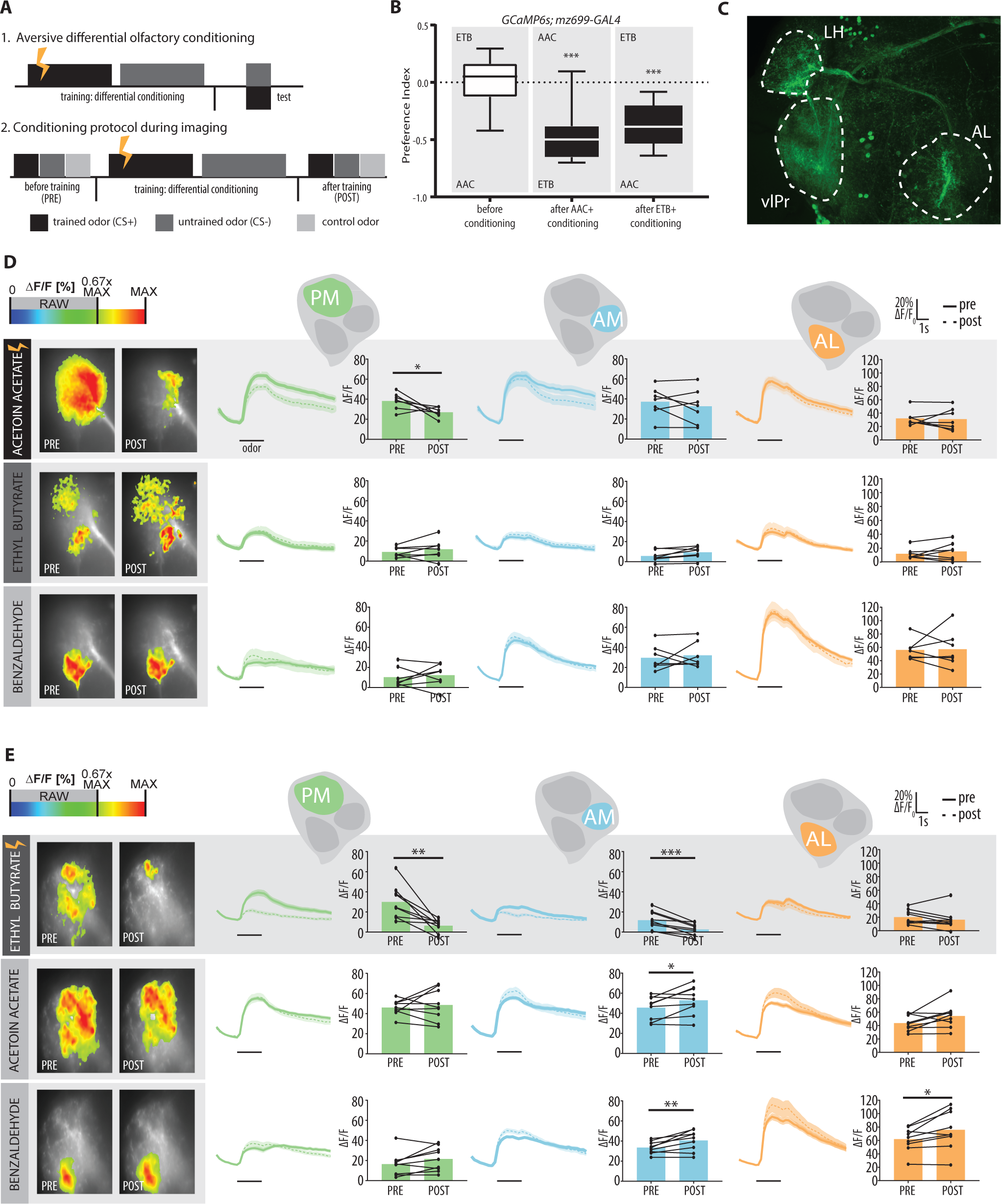
Differential aversive conditioning with attractive odors leads to reduced odor responses in mPNs. (A) Training protocols for differential olfactory learning used in behavioral experiments and functional imaging recordings. (B) Naïve flies are distributed equally between ethyl butyrate (ETB) and acetoin acetate (AAC), but avoid the negative reinforced odor after conditioning (n=16, ***p<0.001, one sample t test against 0; genotype: *MZ699-GAL4; UAS-GCaMP6s*). C) Immunostaining of *MZ699-GAL4* expressing GFP. This line labels mPNs that project from the antennal lobe (AL) to the lateral horn (LH), as well as third-order neurons innervating the LH and ventrolateral protocerebrum (vlPr). (D and E) Functional imaging of flies expressing GCaMP6s under the control of *MZ699-GAL4*. Left: Representative false-color coded images showing odor-induced Ca^2+^ signals before (PRE) and after (POST) differential conditioning for the trained odor (acetoin acetate or ethyl butyrate, respectively) and the control odor (benzaldehyde). Right: time courses of odor-evoked calcium signals for the individual functional LH domain (PM, AM, AL) for each odor tested. Traces represent averaged odor response (±SEM). Black bars represent 2 s odor stimulation. Bar plots give mean activity (ΔF/F) during odor stimulation; individual flies are given by single dots and lines (C: n= 7, D: n=9, paired Student t test, *p<0.05, **p<0.01, ***p<0.001).

We have previously shown that GABAergic mPNs targeting the LH mediate odor attraction (Strutz et al., 2014). Flies unable to release GABA from these neurons no longer prefer previously attractive odorants. Furthermore, within the LH, mPNs responses to odors can be spatially separated into three so-called ‘Odor Response Domains’ (ORDs) where activity correlates with specific odor features, such as odor valence and intensity (Strutz et al., 2014). While the posterior medial domain (LH-PM) shows higher responses to attractive odorants, the anterior lateral domain (LH-AL) responds to aversive odorants in a concentration dependent manner. The third domain, the anterior medial (LH-AL) codes for odor concentration (Strutz et al., 2014). Given the role these neurons play in innate behavior, we asked whether mPNs show experience-dependent changes after aversive conditioning. To monitor the activity of these neurons, we used flies expressing *UAS-GCaMP6s* (Chen et al., 2013) under the control of the enhancer trap line *MZ699-GAL4* (Ito et al., 1997), which labels 86% of mPNs (Lai et al., 2008) (**Figure 1C**). These neurons have their dendritic processes AL and target 75% of all AL glomeruli (Strutz et al., 2014). Most of their synaptic outputs are located in the LH (Liang et al., 2013; Parnas et al., 2013; Shimizu and Stopfer, 2017). Flies were trained under the wide-field microscope and odor-evoked activity was recorded before and immediately after olfactory conditioning. As expected, both attractive odorants, ethyl butyrate and acetoin acetate, but not the aversive odorant benzaldehyde, led to a strong activation in the LH-PM and LH-AM domain (**Figure 1D, E**; **Supplementary Figure 1C**, **D**). Strikingly, associative conditioning reduces the odor-evoked Ca^2+^ activity in the LH-PM domain significantly for the trained odor, in both the absolute and differential conditioning paradigm. This reduction of responses in the LH-PM domain that encodes odor attraction (Strutz et al., 2014), is consistent with learning induced assignment of a negative valence to the innately attractive odor. Interestingly, a slight,but significant increase in the activity in the LH-AM domain was observed for the untrained (CS-) and control odor (which was not presented during the training procedure) in flies differentially trained with ethyl butyrate (**Figure 1E**).

To control for non-associative plasticity in the LH-PM domain, we exposed flies to unpaired presentations of the CS and the US (**Supplementary Figure 2**) or presented each stimulus alone (i.e. odor or electric shock control; **Supplementary Figure 3**). Noteworthy, none of these conditions led to a reduction in the odor-evoked Ca^2+^ activity in the LH-PM domain. It is noteworthy that, in some experiments, responses in the aversive-coding LH-AL for benzaldehyde increased after training (**Figures 1E** and Supplementary Figure 1E). This increase, however, was also present in controls in which the electric shock was delivered alone (**Supplementary Figure 3B**) and might be indicative of a sensitization caused by the electric shock. Taken together, our results show experience-dependent plastiticty in mPNs in the LH including reduced learned odor responses after aversive associative conditioning.

### Plasticity in third-order vlPr neurons

Olfactory input from mPNs is further conveyed to diverse third-order neurons, some of which project from the LH to the ventrolateral protocerebral region (vlPr) (Liang et al., 2013; Strutz et al., 2014). Previous studies have shown that these so-called vlPr neurons are inhibited by mPNs in the LH in an odor-dependent manner. Thus, we wondered whether the learning-induced plasticity in the mPNs is reflected in vlPr neuron activity. To examine this question we used the line *VT063948-GAL4* (Tirian and Dickson, 2017) to express the calcium sensor GCaMP6s in vlPr neurons. This enhancer line also labels neurons outside of the vlPr, but we restricted the imaging window to the ventral area of the LH, where the highest overlap with MZ699+ neurons is expected.

Previous studies have shown that vlPr neurons (also called AV1 tract neurons in some studies) preferentially respond to aversive olfactory stimuli (Huoviala et al., 2020; Liang et al., 2013; Mohamed et al., 2019; Strutz et al., 2014). In our experiments we find that the highest response was indeed induced by the aversive odorant benzaldehyde. However, the neurons responded also, although less strong, to the attractive odors acetoin acetate and ethyl butyrate (**Figure 2**). Again, odor-evoked responses were measured before and after differential olfactory conditioning. Indeed, the responses for the trainined odor after conditioing was significantly increased (**Figure 2B**). This increase was training specific, as neither the shock nor olfactory stimulation alone generated a difference in the calcium signals before and after conditioning (**Figure 2C, D**).

**Figure 2.**
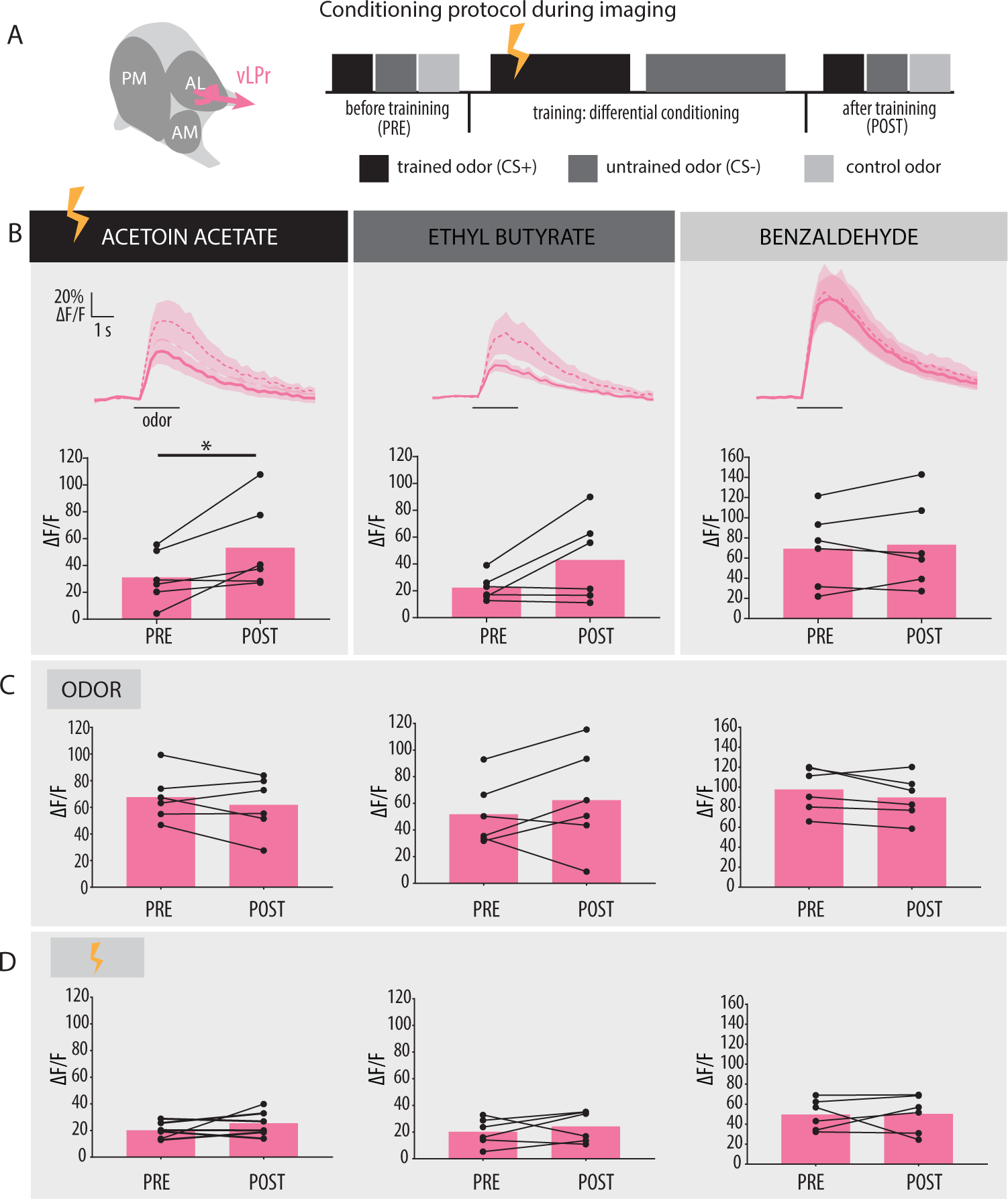
Aversive differential olfactory conditioning leads to increased odor responses in third-order vlPr neurons after training. (A) Functional imaging of flies expressing *UAS-GCaMP6s* in a subset of vlPr neurons using *VT063948-GAL4* before (PRE) and after (POST) aversive olfactory conditioning. (B) Odor-evoked responses show a significant increase for the trained odor (acetoin acetate) after conditioning (n=6, *p<0.05). Top, odor-evoked calcium responses for each odor averaged over all flies (±SEM). Black bars indicate the odor delivery for 2 s. Bottom, bar plots give mean activity upon odor stimulation; individual flies are given by single dots and lines. No significant difference was observed in mock-trained flies in which (C) the odor was presented without negative reinforcement (n=6, ns) or (D) the electric shock was applied in the absence of olfactory stimuli (n=6, ns). All statistics are paired Student t test.

### Aversive conditioning affects odor-evoked responses of mPNs in the antennal lobe

While we know that mPNs have most of their outputs in the LH (Parnas et al., 2013), with a small number of recurring processes in the AL (Okada et al., 2009; Shimizu and Stopfer, 2017), most of mPN postsynapses are located in the AL, where they gather information across glomeruli. We thus hypothesized that changes in activity in mPNs could be present already in this first olfactory processing center, i.e. at their postsynaptic dendrites and not only at their axonal target regions in the LH. Imaging from mPNs in the AL is challenging, as dendritic processes are sparse and diffuse (Strutz et al., 2014). To overcome this difficulty, we used flies co-expressing *tdTotmato* in uniglomerular excitatory PNs (i.e. *GH146-QF*) to visualize glomerular borders, and *UAS-GCaMP6s* expression in the *MZ699-GAL4* enhancer trap line to monitor odor responses in mPNs (**Figure 3A**). This approach allowed us to identify glomeruli and adjust the focal plane, based on the fluorescence from uniglomerular neurons. Flies were trained using an absolute conditioning protocol and odor-evoked Ca^2+^ responses were quantified from those glomeruli we could reliably identify among all flies measured (**Figure 3B**). Our results revealed a significant decrease in GCaMP6s fluorescence intensity for the trained odor in all, but one (i.e. DM5), of the identified glomeruli (**Figure 3C**), after associative conditioning. Interestingly, DM5 was the only glomerulus to show a significant decrease after conditioning to the aversive control odorant benzaldehyde. These results show that the decrease in odor-evoked responses of mPNs observed after aversive conditioning occurs in the AL indicating that experience-dependent changes are established already at the first olfactory processing level.

**Figure 3.**
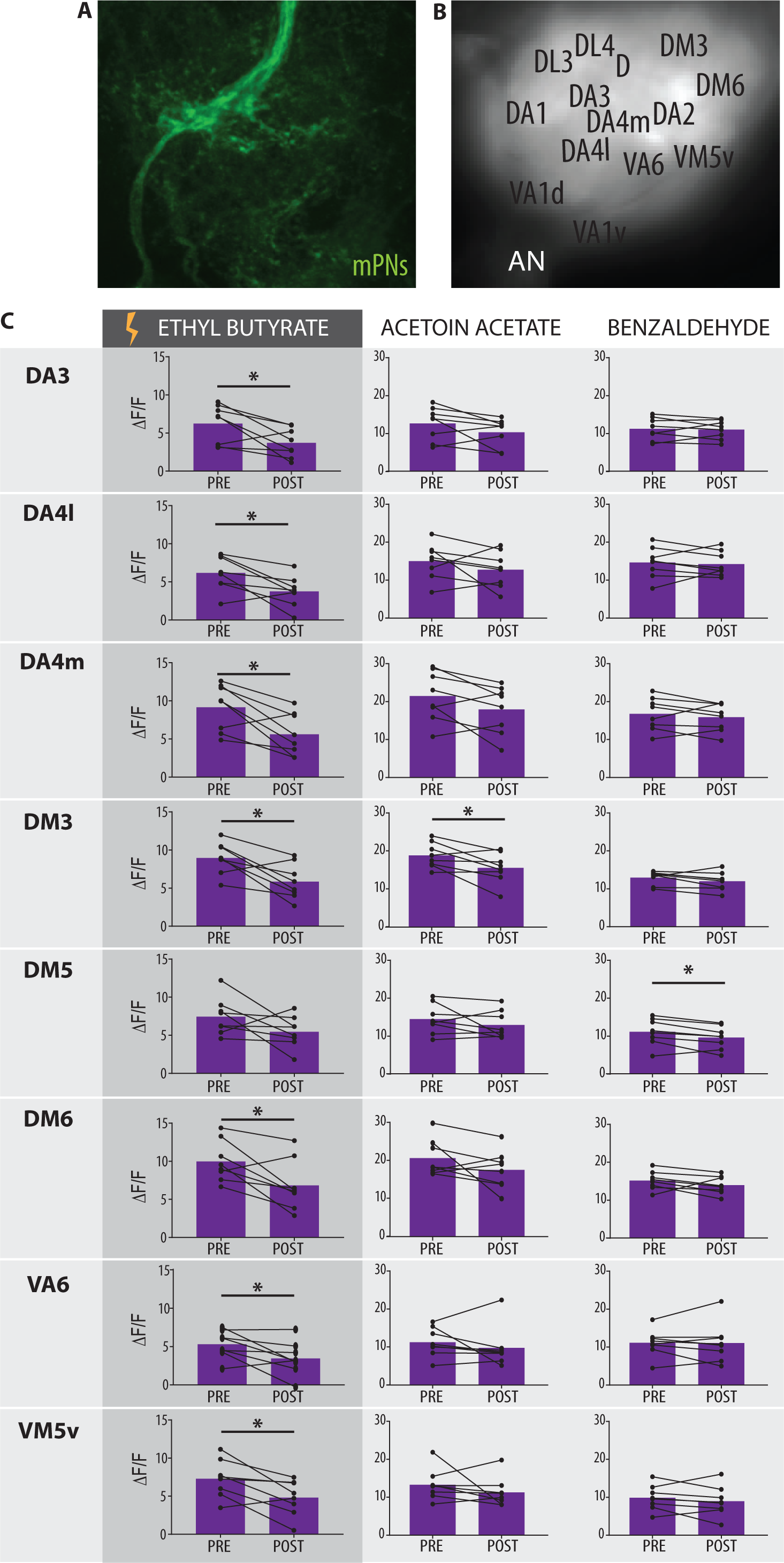
Aversive differential conditioning leads to decreased odor responses in dendrites of mPNs in the antennal lobe for the trained odor. (A) Immuno-staining showing innervation of MZ699+ neurons in the antennal lobe (AL) (*MZ699-GAL4;UAS-GCaMP6s*, antibody against GFP). (B) Flies co-expressing tdTomato in GH146+ neurons (*GH146-QF; QUAS-tdTomato*) and GCaMP6s in MZ699+ neurons were used in calcium imaging experiments. Fluorescence from GH146+ neurons was used for the identification of a subgroup of glomeruli. (C) Functional Ca^2+^imaging of flies expressing GCaMP6s under the control of *MZ699-GAL4*. Bar plots give mean activity during odor stimulation in individual glomeruli before (PRE) and after (POST) absolute aversive conditioning for the trained odor (ethyl butyrate) and control odors (acetoin acetate, benzaldehyde); individual flies are given by single dots and lines (n= 8, paired Student t test, *p<0.05).

### Glutamate and serotonin modulate odor responses in mPNs after aversive conditioning

We showed that learning induced plasiticty in mPNs. Given that the changes in trained-odor response are detected at the level of the AL, it is plausible that plastiticty is driven by AL-intrinsic modulatory pathways. Modulatory neurons are widespread in the AL, with local interneurons (LNs) innervating several or all glomeruli (Chou et al., 2010; Wilson, 2013). Among them, GABA-immunoreactive LNs represent the predominant type (Chou et al., 2010). Thus, we first investigated whether GABAergic input is required for the modulation of mPNs, by expressing *UAS-Rdl RNAi* against the Rdl subunit of the GABA_A_ receptor under the control of *MZ699-GAL4*. However, flies in which this receptor was downregulated still revealed a modulated odor response in mPNs following aversive conditioning (**Figure 4A**). In addition to GABA, a subpopulation of LNs is glutamatergic whose ventral position of their cell bodies in the AL coincides with that of MZ699+ mPNs, and both neuronal types projecting to the AL through the ventral AL fascicle (Chou et al., 2010; Das et al., 2011; Sizemore and Dacks, 2016). Furthermore, glutamate has been shown to have an inhibitory role, mediated by the ionotropic GluClα receptor subunit (Liu and Wilson, 2013), in AL processing. We therefore employed *UAS-GluClα-RNAi* to knock-down the expression of GluClα in mPNs (**Figure 4B**). Notably, in flies expressing the UAS*-GluClα-RNAi* transgene the learning induced plasticity in mPNs was abolished. Instead, we observed a significant increase in Ca^2+^ evoked activity in response to the trained odor in the LH-PM domain, but also in the LH-AM and LH-AL domains (**Supplementary Figure 4**). These changes, however, were specific to the trained odor, and were absent in the control odorants. Our results thus suggest that glutamate signaling, which seem to have an inhibitory impact on mPNs, is required to modulate odor-evoked mPN responses after aversive training.

**Figure 4.**
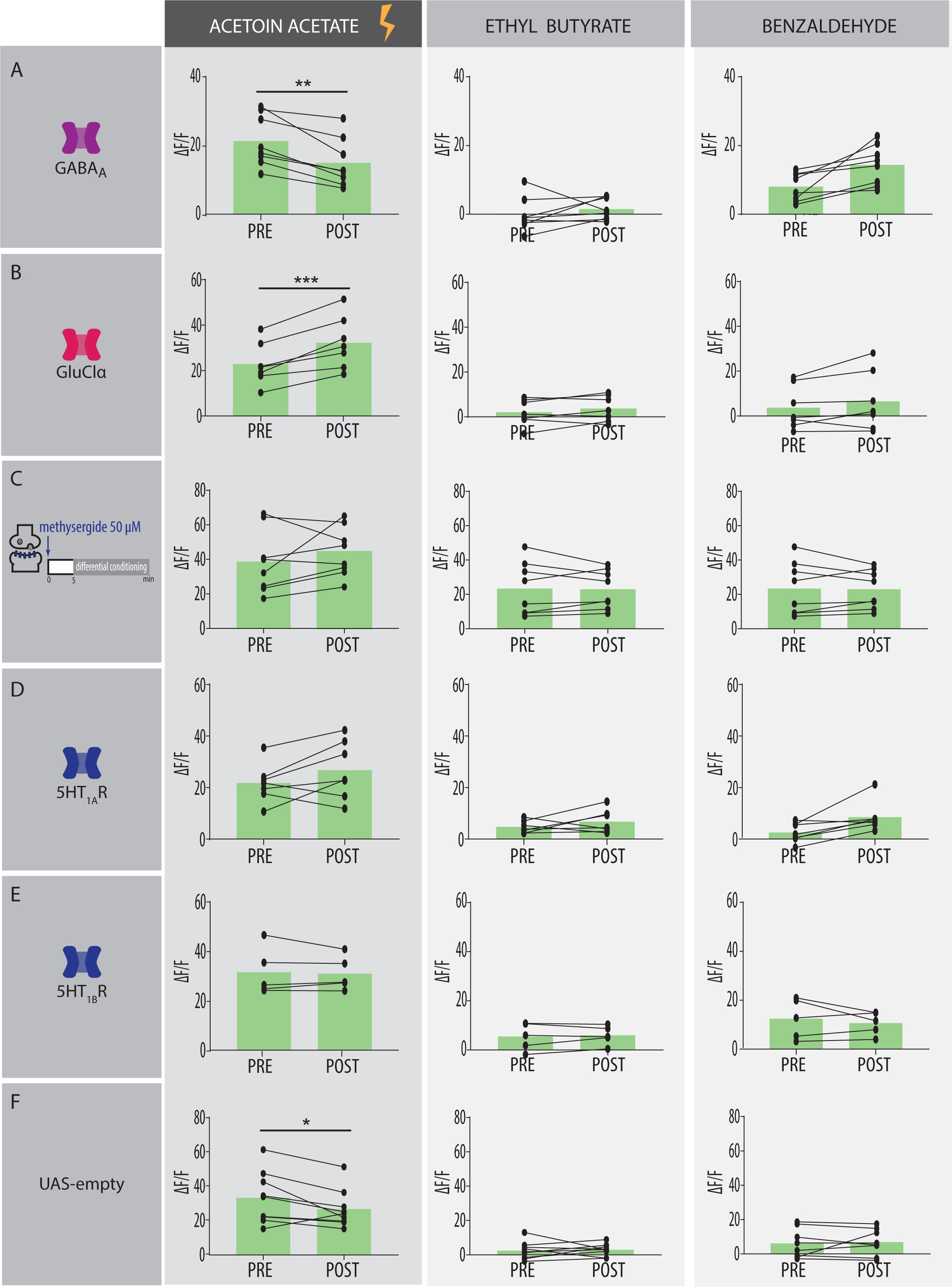
Learning-induced response decrease in mPNs after training requires glutamate and serotonin receptor activity. Odor-evoked peak responses in mPNs before (PRE) and after (POST) aversive differential conditioning, from flies presenting the genetic or pharmacological manipulation indicated on the left. Acetoin acetate was used as the trained odor, while ethyl butyrate and benzaldehyde served as control odors. (A) Expression of *UAS-RdliG-GABA_A_ RNAi* was used to downregulate GABA-A receptor expression in MZ699+ neurons. Odor-evoked responses to the trained odor are significantly decreased after training in mPNs, showing that GABA-A receptor downregulation does not affect the training-induced response decrease (n=8, **p<0.01). (B) Downregulation of glutamate receptors in MZ699+ neurons via expression of *UAS-GluClα RNAi* reverses changes in the odor-evoked activity of mPNs after conditioning (n=7, ***p<0.001). (C) Pharmacologically inactivating serotonin (5HT) receptors using the receptor antagonist methysergide abolishes learning-induced reduction of the odor responses in mPNs (n=8, ns, paired Student t test). (D, E) Downregulating inhibitory 5HT receptor subunits using either *UAS-5HT1A R*NAi or *UAS-5HT1B RNAi* in MZ699+ neurons prevents changes in odor responses of mPNs after conditioning (5HT1A: n=5; 5HT1B: n=7, ns). (F) Flies expressing the control transgenic construct *UAS-empty RNAi* in MZ699+ neurons show significantly reduced odor response in mPNs after conditioning (n=8, *p<0.05). All statistics are paired Student t test. Peak fluorescence is shown for the PM functional LH domain (also see Figure S5).

In addition to the inhibitory neurotransmitters GABA and glutamate also serotonin has been shown to be released in the AL and to modulate the activity of PNs (Dacks et al., 2009; Zhang and Gaudry, 2016). In *Drosophila*, in-situ hybridization experiments have revealed five types of serotonin receptors (5HTR) in ventral PNs (Sizemore and Dacks, 2016), a broader group to which MZ699+ mPNs belong to. To address the role of serotonin in mPNs, we started by applying 50µM methysergide, an antagonist of 5HT receptors, five minutes before the onset of differential conditioning. Notably, pharmacologically blocking serotonin receptors abolished the decrease of the odor response in mPNs that was observed after conditioning in our previous experiments (**Figure 4C**). In a second step, we assessed the involvement of specific 5HT receptor subunits. Given our results, we presumed that 5HT could have a modulatory role on mPNs. The role of serotonin in the AL depends greatly on the specific 5HT receptors that the cells express, with two of the five serotonin receptor subunits mediating inhibition. In adult flies, these correspond to 5HT1A and 5HT1B receptor subunits (Sizemore and Dacks, 2016). We expressed RNAi against these subunits and examined whether changes in mPNs after aversive conditioning still occurred. Downregulating of either 5HT1A or 5HT1B in mPNs prevented any modulation of mPN odor responses after aversive training from taking place (**Figure 4D, E**). Expression of *UAS-empty-RNAi* served as a control and showed a significant response decrease for the trained odor after conditioning (**Figure 4F**). Taken together, our results indicate that the function of inhibitory glutamate and serotonin receptor subunits are required for the changes in activity observed in mPNs following aversive associative learning.

### Activity from mPNs is not required for aversive olfactory conditioning

Previous work showed that downregulation of GABA release from mPNs leads to a reduction in attraction to innately attractive odorants (Strutz et al., 2014). Here, we have shown that flies can learn to avoid an attractive odorant and that aversive olfactory conditioning results in a decrease in the activity of mPNs, possibly associated to a decrease in GABA release from their synaptic terminals. Therefore, we hypothesized that preventing modulation from taking place in mPNs might result in learning deficits in flies. Using RNAi transgenes, we independently downregulated the expression of glutamate and serotonin receptor subunits selectively in MZ699+ neurons. Unexpectedly, after aversive conditioning, none of the experimental manipulations resulted in a learning deficit, compared to the genetic controls (**Supplementary Figure 5A-C**). We additionally evaluated the role of the excitatory serotonin subunit 5HT7 in aversive conditioning experiments, as a recent study proposed its involvement in PN modulation (Vogt et al., 2020), but also did not observe any effect on the learning performance when this subunit was downregulated selectively in MZ699+ neurons (**Supplementary Figure 5D**).

In principle, individually downregulating receptor subunits might not result in a behavioral phenotype as the remaining subunits might compensate for any deficits. In addition, RNAi-mediated downregulation is rarely complete. Therefore, we sought to manipulate serotonin input to mPNs by artificially preventing its release from the modulatory neurons. The contralaterally-projecting serotonin immunoreactive deuterocerebral neuron (CSDn), innervating first and second order olfactory neuropils, has been proposed as the major source of serotonin in the AL (Dacks et al., 2009; Zhang and Gaudry, 2016). CSDn have been shown to modulate appetitive and pheromone olfactory responses in the AL (Xu, 2016; Singh, 2013), but its role in associative learning has, to our knowledge, never been assessed. We expressed the thermo-sensitive transgene *shibire* specifically in CSDns (R60F02-GAL4) to block synaptic vesicle transmission from these neurons. Our results show that blocking CSDn synaptic output does not impair associative learning (**Supplementary Figure 5E**). Taken together, these results suggest that glutamate and serotonin release onto mPNs is involved in modulating odor responses of mPNs after learning, but seem not to be required for the behavioral expression of the aversive associative memory.

Finally, we investigated the role of mPNs regarding innate and learned behavior by analyzing whether the output of MZ699+ PNs is required for memory formation following aversive olfactory conditioning. However, blocking synaptic release from MZ699+ neurons using the temperature-sensitive *shibire* did not affect the innate preference between acetoin acetate and ethyl butyrate in untrained flies nor did it result in any learning deficit in differential conditioning experiments, compared to control flies (**Figure 5A**). As blocking synaptic output from MZ699+ neurons might also impair the internal representation of the unpunished attractive odorant in differential conditioning (i.e. CS-), making it difficult to see a clear behavioral phenotype, we opted for absolute conditioning. When an attractive odor was presented against a solvent, flies whose MZ699+ output was impaired showed a reduced preference for the attractive odorant (**Figure 5B**), confirming previous data (Strutz et al., 2014). Noteworthy, this avoidance was further enhanced after absolute conditioning, compared to genetic controls, and absent in temperature controls (**Figure 5C**). However, when flies were trained to one odor at the restrictive temperature of 32°C and were immediately tested at the permissive temperature of 23°C against the trained and a novel attractive odor, no difference in learning was observed between experimental and control flies (**Figure 5D**). These findings show that output from mPNs is dispensable for associating an aversive event with an appetitive odor.

**Figure 5.**
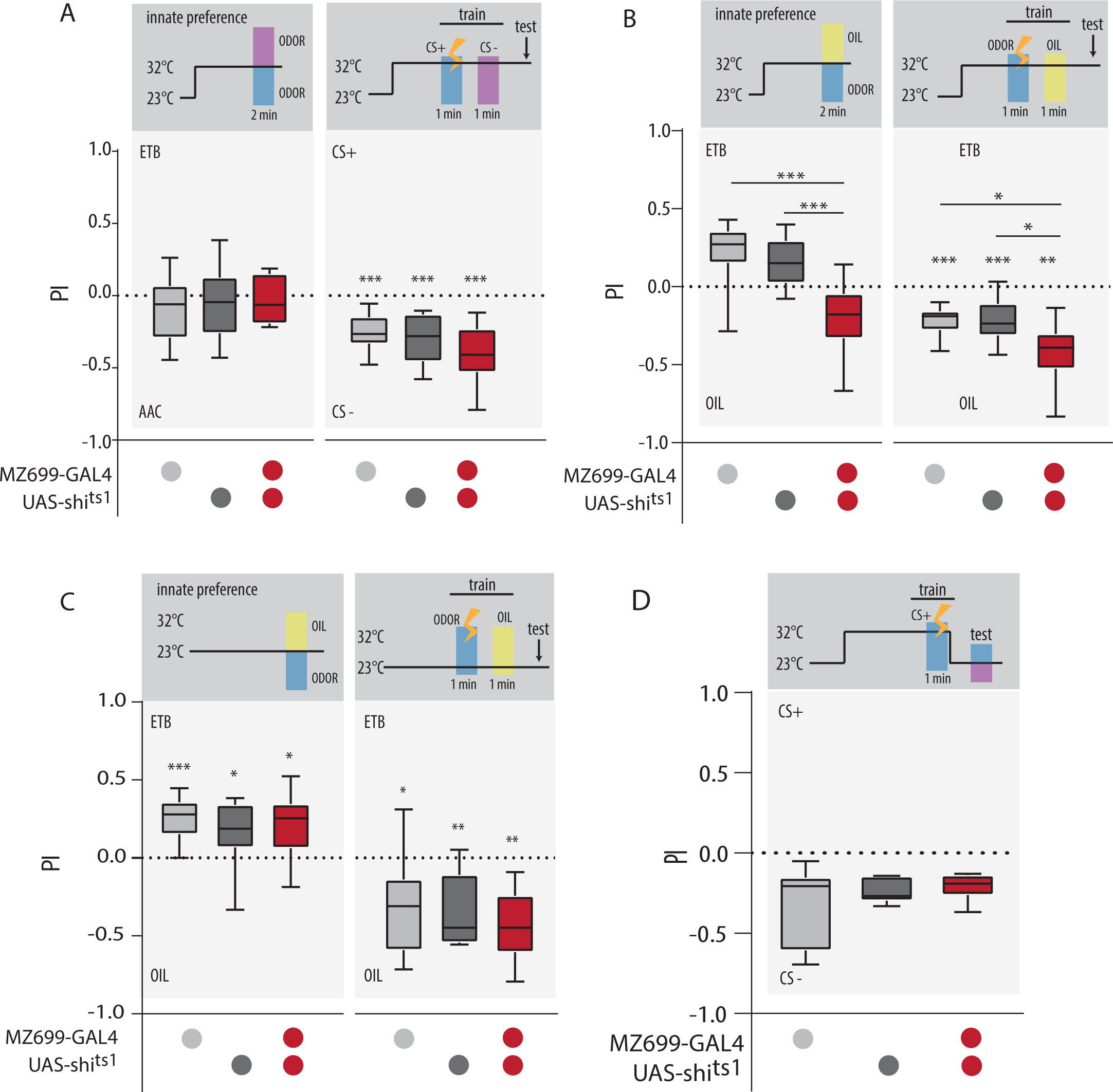
Blocking MZ699+ neurons does not impair aversive olfactory associative learning. (A) Left, innate preference between acetoin acetate and ethyl butyrate at the restrictive temperature 32°C in flies expressing *UAS-shibire^ts1^* under the control of *MZ699-GAL4*, and the corresponding genetic controls (n= 11-13, ns). Right, preference for the trained (CS+) vs. untrained (CS-) odor after differential aversive conditioning at the restrictive temperature 32°C for the same fly lines (n=11-13, ***p<0.001) (B) Left, innate preference between ethyl butyrate and paraffin oil at the restrictive temperature 32°C in flies expressing *UAS-shibire^ts1^* under the control of *MZ699-GAL4*, and the corresponding genetic controls (n= 12-16, ***p<0.001). Right, preference for the trained odor ethyl butyrate vs. paraffin oil after absolute aversive conditioning at the restrictive temperature 32°C for the same fly lines (n= 14-17, *p<0.05, **p<0.01, ***p<0.001). (C) Innate preference for ethyl butyrate against paraffin oil (left, n=10, ns) and after absolute aversive conditioning (right, n=10, ns) at the permissive temperature 23°C for flies expressing *UAS-shibire^ts1^* under the control of *MZ699-GAL4*, and in the corresponding genetic controls. Asterisks denote significant differences between groups (one-way ANOVA followed by a Tukey’s multiple comparisons post-hoc test) or a significant difference against 0 (one sample Student t test). (D) Top, scheme illustrating training protocol for differential aversive olfactory conditioning with ethyl butyrate or acetoin acetate as CS+, respectively. The output of MZ699+ neurons was silenced during training using expression of *shibire^ts1^* at the restrictive temperature (i.e. 32°C), while flies were tested at the permissive temperature (i.e. 23°C) when mPNs were functional. Bottom, silencing the output of MZ699+ neurons during learning did not result in preference differences during testing compared to genetic controls (n= 9-10, one-way ANOVA followed by Tukey’s multiple comparisons post-hoc test, ns).

### Appetitive learning modulates mPNs

Since activity in mPNs has been shown to mediate odor attraction, we next asked whether pairing an odor with a reward would result in an increase in the activity of these neurons. In order to examine whether mPNs are bidirectionally modulated, we utilized the sugar -based appetitive conditioning paradigm in flies. Flies expressing *GCaMP6s* under the control of *MZ699-GAL4* were exposed to a 2 min. odor stimulation paired with sucrose (**Figure 6**). Trained flies were dissected and tested 3 hours later under the microscope, where we monitored odor-evoked responses in mPNs. A group receiving only sucrose for 2 min. was used as experimental control. Strikingly, trained flies showed a significant increase in Ca^2+^ activity in the ‘attraction-coding’ PM-LH domain to the trained odor. Together with our previous findings, these results imply that the modulation of mPNs is bidirectional and specific to the trained odor used. Taken together, mPN activity is thus reduced in aversive conditioning, and increased by appetitive learning.

**Figure 6.**
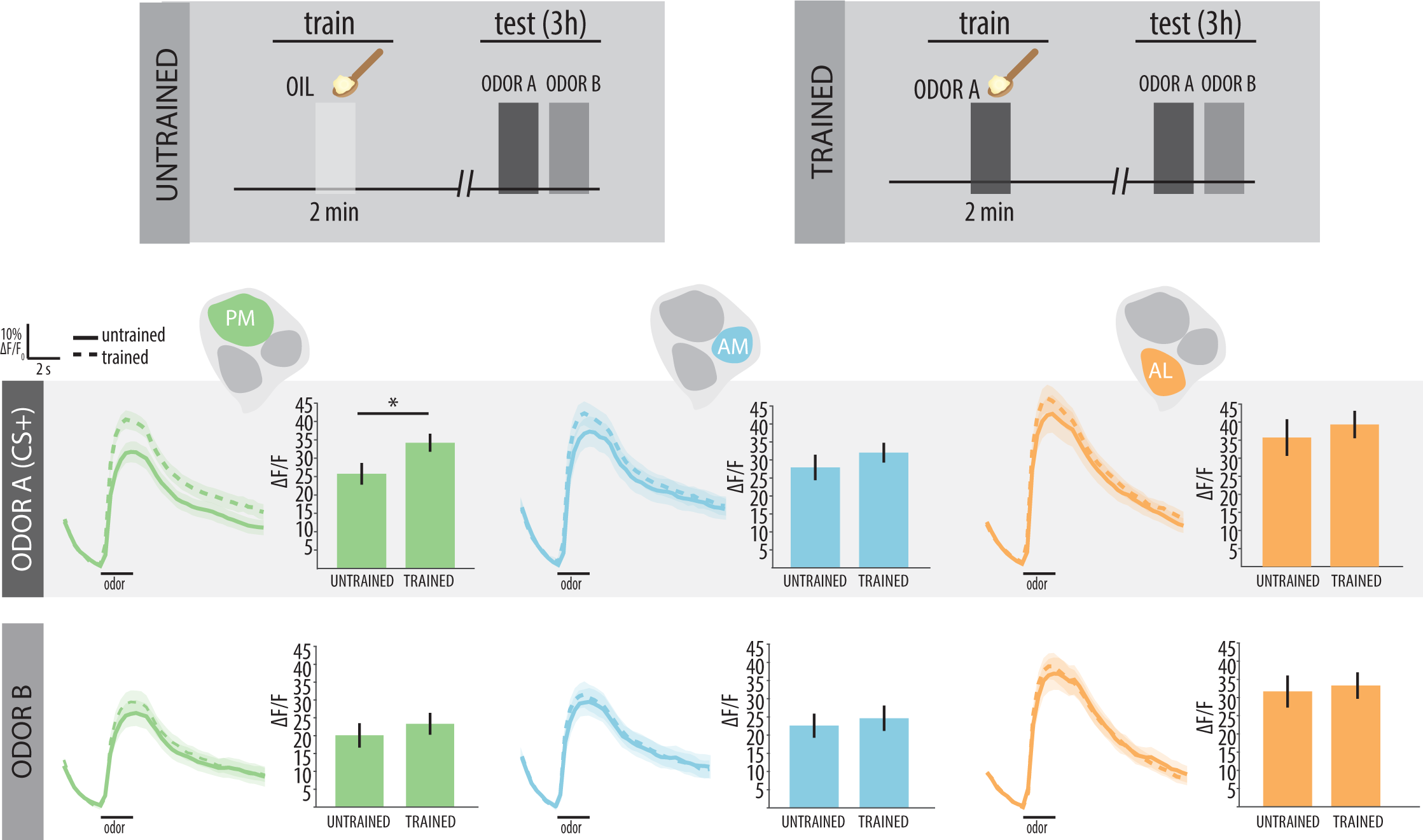
Appetitive olfactory conditioning enhances odor-evoked CS+ responses in mPNs. Top: diagrams illustrate conditionings protocols. Flies expressing GCaMP6s in mPNs using *MZ699-GAL4*were trained by pairing a 2 min odor presentation with a saturated sucrose reward, and tested 3 hr later under the microscope (i.e. trained). As trained odorants, acetoin acetate and ethyl butyrate were alternatively used as CS+. In the control group, flies received the sugar reward without odor presentation (i.e. untrained). For both groups, during imaging, responses were measured for the trained odor (i.e. odor A) and an additional control odor (i.e. odor B). When acetoin acetate was used as CS+, ethyl butyrate was used as control, and vice versa. Bottom: time traces of Ca^2+^ signals averaged over all flies for the CS+ (i.e. ODOR A) and the control odor (i.e. ODOR B) (±SEM) for each functional LH domain (PM, AM, AL) and each odor tested, given for trained and untrained groups. Bar plots give mean activity after odor stimulation (n= 13-19; unpaired Student t test; *p<0.05).

### mPNs are required in the retrieval of appetitive memories

We next determined whether the output of the mPNs is required in appetitive olfactory conditioning with sucrose reinforcement. To do so, flies expressing *shibire^ts1^* in MZ699+ neurons were trained to associate an odor with a sugar reward, at the permissive temperature of 23°C. In order to assess the involvement of MZ699+ neurons in memory retrieval, we blocked their output by raising the temperature to 30°C (i.e. restrictive temperature) 30 min before and during testing of 3 h appetitive memory. Strikingly, silencing MZ699+ neurons led to a significant memory impairment (**Figure 7A**). Importantly, no memory deficit was observed in genetic controls expressing *MZ699-GAL4* or *UAS-shi^ts1^* (**Figure 7A**) or in flies tested at the permissive temperature of 23°C (**Supplementary Figure 6)**. Our results thus demonstrate that mPNs are required for the retrieval of appetitive olfactory memories.

**Figure 7.**
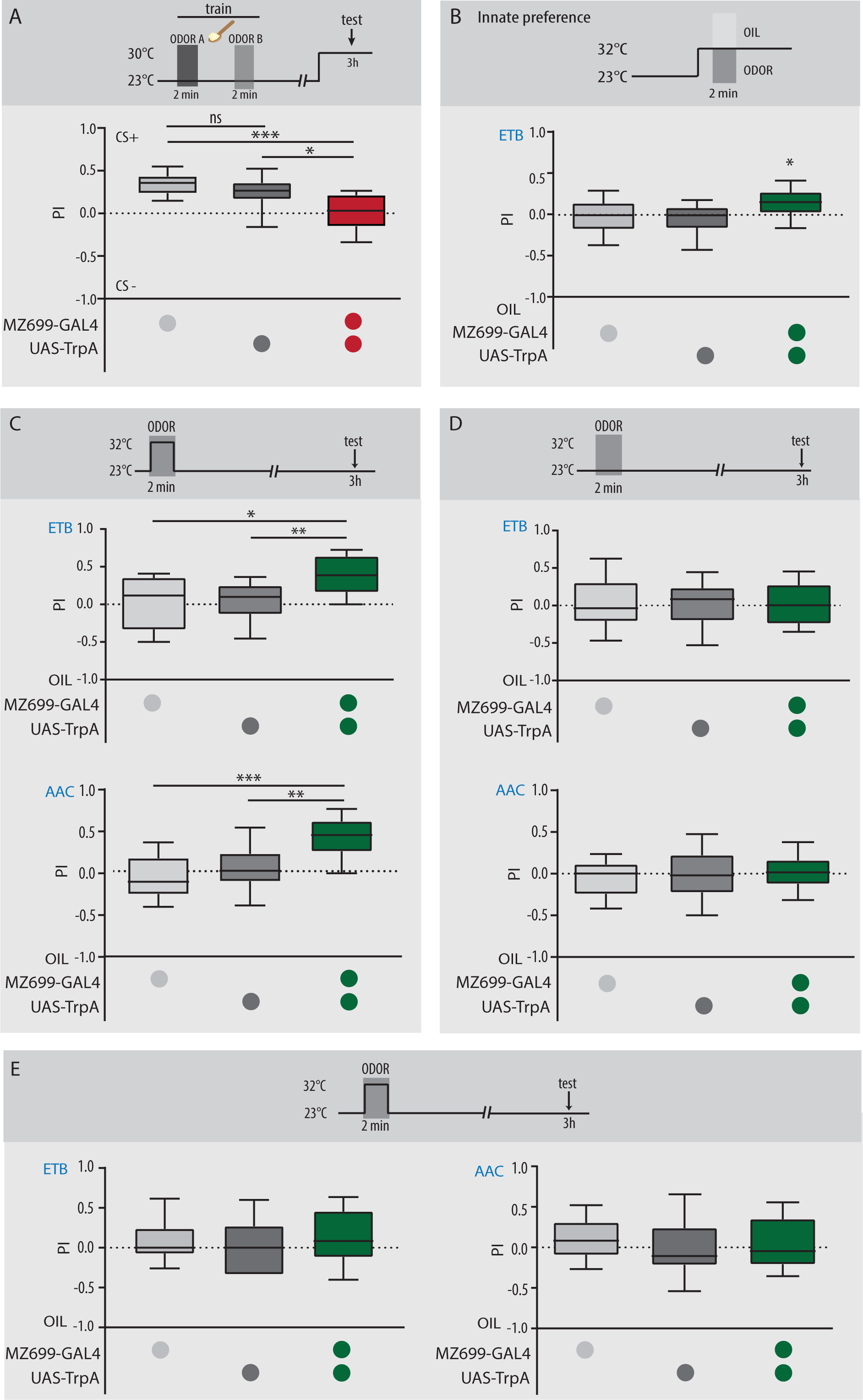
MZ699+ neurons are required for the retrieval of appetitive memories and are sufficient to generate an appetitive memory. (A) Top, scheme illustrating training protocol for differential appetitive olfactory conditioning with 3-octanol or 4-methylcyclohexanol as CS+. Flies were trained at the permissive temperature (i.e. 23°C) and tested 3 h later at the restrictive temperature of 30°C. Bottom: Flies expressing *shibire^ts1^* show impaired learning performance compared to genetic controls (n= 12, one-way ANOVA with Tukey’s multiple comparisons post-hoc test, *p<0.05, ***p<0.001). (B) Flies expressing *TrpA1* under the control of *MZ699-GAL4* show a significant preference for ethyl butyrate which is absent in genetic controls (n=8-10, one-sample t-test against 0, #p<0.05). No significant difference is observed between the groups (one-way ANOVA). (C) Top, scheme illustrating artificial conditioning protocol used in flies expressing *TrpA1* under the control of *MZ699-GAL4*. Bottom, flies expressing *TrpA1* significantly prefer the odor that was presented during heat activation (i.e. ethyl butyrate or acetoin acetate, respectively), compared to the genetic controls, when tested 3 h later at 23°C (n= 10-12 for ethyl butyrate, n= 12 for acetoin acetate, *p<0.05, **p<0.01, ***p<0.001). (D) Top, scheme illustrating control training protocol, with odor stimulation at 23°C. Bottom, no significant difference in odor preference is observed between experimental lines and genetic controls at 23°C (n= 12 for ethyl butyrate and acetoin acetate, ns). All statistics are one-way ANOVA with Tukey’s multiple comparisons post-hoc test. (E) Top, scheme illustrating control training protocol used in satiated flies expressing *TrpA1* under the control of *MZ699-GAL4*. Bottom, no significant difference in odor preference is observed between experimental lines and genetic controls (n=10-12 for ethyl butyrate and n=11-12 for acetoin acetate, ns). All statistics are one-way ANOVA with Tukey’s multiple comparisons post-hoc test.

### Artificial activation of mPNs generates appetitive memories

Next, since mPNs are required for appetitive learning, we next wondered whether artificially activating these neurons during odor presentation would lead to the formation of appetitive memories. Expression of the transgene for the cation thermo-sensitive channel *TrpA1* allows temporal control of neuronal activation by exposing flies to temperatures over 25°C (Hamada et al., 2008). In our experiments, we restricted the expression of *TrpA1* to MZ699+ neurons, and flies carrying the transgenes *UAS-TrpA1*, *MZ699-GAL4* or *MZ699-GAL4*; *TrpA1* were submitted to a 2-choice assay between ethyl butyrate and paraffin oil at the activating temperature of 32°C. In these experiments, we used ethyl butyrate at a concentration at which flies showed no innate preference. Interestingly, artificially activating MZ699+ neurons was sufficient to drive odor attraction, as neither *TrpA1* nor *GAL4* controls showed any attraction towards the odor (**Figure 7B**). We then wondered whether pairing an odor with the artificial activation of these neurons would be sufficient to implant an appetitive memory in flies. Thus, starved *UAS-TrpA1*, *MZ699-GAL4* and *MZ699-GAL4*;*UAS-TrpA1* flies were conditioned by presenting an odor at the activating temperature. All groups were tested 3 hr later at 23°C. Surprisingly, *MZ699-GAL4;UAS-TrpA* flies exhibited a robust appetitive memory that was statistically different from all control groups (**Figure 7C**). Notably, this memory was absent in temperature controls (**Figure 7D**) or ad-libitum fed flies (**Figure 7E)**. In sum, our results show that not only artificial activation of MZ699+ mPNs drives approach, but that depolarizing these neurons during odor presentation is sufficient to implant appetitive memories in the fly’s brain.

## Discussion

In both vertebrate and invertebrate systems, the question of how experience modifies innate behavior and its underlying circuits remains largely unanswered. In flies, several studies showed that neuronal circuits involved in learning also mediate innate responses (Aso et al., 2014; Dolan et al., 2018; Kadow, 2019), very few assessed whether innate circuits show experience dependent-modulation or are required for olfactory learning (Dolan et al., 2018; Séjourné et al., 2011). In this study we took advantage of the concise olfactory circuit in *Drosophila* to examine whether olfactory circuits involved in innate behavioral responses are modulated by and required for associative olfactory learning.

### mPNs are bidirectionally modulated after associative conditioning

In behavioral and imaging experiments, we demonstrate that olfactory associative learning modulates the activity of mPNs as summarized in a circuit model (**Figure 8**). While olfactory aversive conditioning results in a decrease in the activity of these neurons, appetitive conditioning leads to an increase of their activity. These changes are correlated with behavior, where silencing mPN synaptic output drives odor avoidance, while activation promotes odor attraction. Bidirectional modulation of brain circuits has been previously observed both in *Drosophila* (Handler et al., 2019; Owald et al., 2015; Yamagata et al., 2016) and mammals (Huang et al., 2004; Seamans et al., 2001; Shen et al., 2008), in the context of learned behavior. Our results suggest that a similar principle may underlie modulation of circuits specifically involved in innate olfactory responses. In previous studies, mPNs have been shown to be required in approach behavior (Strutz et al., 2014). Here we show that associative learning results in the modulation of these neurons, driving or suppressing approach behavior according to the value of the CS.

**Figure 8.**
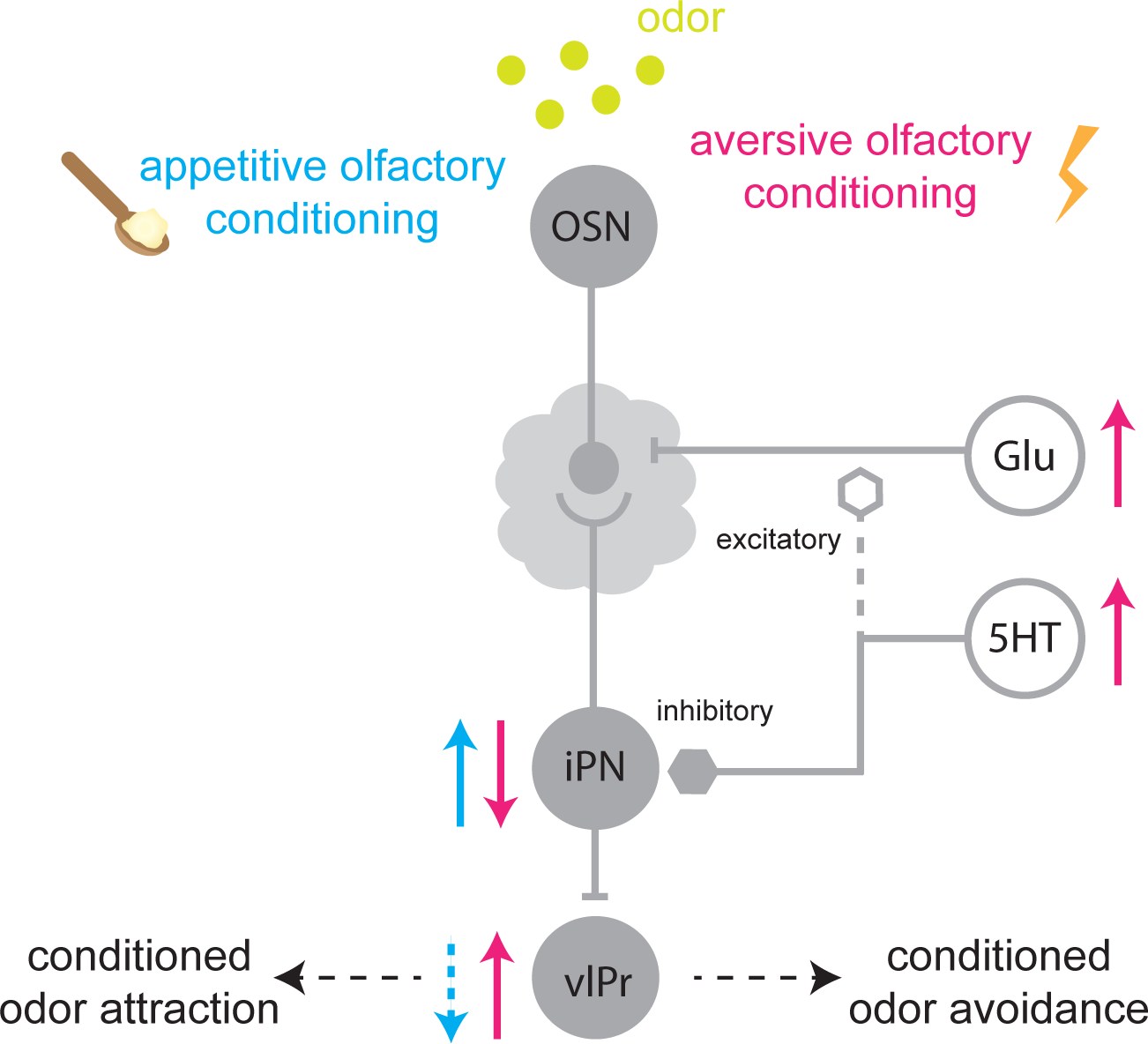
Model for associative learning involving mPNs. Odors are detected by olfactory sensory neurons (OSN) that synapse onto inhibitory, multiglomerular projection neurons (mPN). During aversive olfactory conditioning, increased glutamate (Glu) and serotonin (5HT) release leads to a decrease in the activity of mPNs, which results in an activity increase in downstream ventrolateral protecerebral region (vlPr) neurons. Appetitive olfactory conditioning leads to an increase in the activity of mPNs, whose putative increase of GABA release at their synaptic terminals might result in a decreased activity in vlPr neurons.

In calcium imaging experiments we reveal that the modulation of odor-evoked responses in mPNs is already present in the AL, where these neurons receive their synaptic input. There, mPNs are in close proximity to local and modulatory neurons, whose activity may modify odor-evoked responses in mPNs. In particular, glutamate and serotonin have been shown to be involved in processing of olfactory information in flies and other invertebrates (Dacks et al. 2009; Sizemore and Dacks 2016; Zhang and Gaudry 2016; Xia et al. 2005; Troncoso and Maldonado 2002; Burrell and Sahley 2004). However, the role of these neurotransmitters in the modulation of olfactory information during learning remains to be established. Here we show that serotoninergic and glutamatergic input account for the mPN modulation after aversive olfactory conditioning. Downregulating the ionotropic receptor subunit GluClα reverses the reduction of mPN activity after conditioning. Similarly, blocking the inhibitory subunits of serotonin receptors prevents mPNs from showing activity changes after conditioning. These observations suggest a role for these neurotransmitters in fine tuning the activity of olfactory circuits that is activity-dependent. In behavioral experiments, however, independently downregulating each receptor subunit did not result in learning deficits. Several lines of reasoning may account for this. First, RNAi downregulation is rarely complete (McManus et al., 2002), and remaining receptor expression might be sufficient to produce behavior that is indistinguishable from controls. Second, the coordinated activity of more than one subunit might be required to sustain a learning process; downregulating one subunit at a time might not be sufficient. Third, during our experiments, other neural circuits involved in olfactory learning (e.g. neurons in the MBs) are intact. Their activity may override any possible changes or deficits taking place in parallel circuits. It is also possible that changes in mPNs, mediated by these receptor subunits, actually do not have an impact on olfactory learning. However, binary decision assays, like the one used here, do not capture the spatial and temporal dynamics of decision-making. This robust, but broad, behavioral paradigm might not reflect the exquisite subtleties of fine olfactory tuning.

Similarly, transiently blocking mPN output during aversive olfactory learning did not result in a learning impairment. In light of our imaging data, this result is not surprising. As mentioned above, aversive olfactory conditioning results in a decrease in the activity of mPNs. Probably, artificially silencing these neurons contributes even further to this activity reduction. Interestingly, in both mammals and invertebrates a reduction in inhibition has been shown to take place during acquisition and retrieval of aversive memories (Letzkus et al., 2015; Liu and Davis, 2009; Zhou et al., 2019). In *Drosophila*, the GABAergic anterior paired LH neuron (APL) is suppressed by olfactory learning (Liu and Davis, 2009), and knocking down the GABA_A_ receptor RDL in neurons postsynaptic to APL enhances aversive olfactory learning (Liu et al., 2007). Although in our experiments we cannot conclude that silencing mPNs results in an enhancement of olfactory memory, a recent study did show the involvement of mPNs in the enhancement of a 24h context dependent olfactory memory. This suggests that mPN activity might impact behavior in situations where more than one sensory modality is involved. This goes in line with our data from LH output neurons (i.e. vlPr neurons): In calcium imaging experiments, we show that the activity of these vlPr neurons, downstream to mPNs, increases after aversive conditioning. Noteworthy, this is likely due to a decrease in the inhibitory input from mPNs onto vlPr neurons, and most likely not from changes in the activity of excitatory uniglomerular PNs that target the same region (Liang et al., 2013)). Given our current understanding of the LH circuitry, it is highly likely that the vlPr neurons studied here receive input from other sensory modalities and integrate olfactory information in the framework of additional available sensory information (Chin et al., 2018; Huoviala et al., 2018; Marin et al., 2020). Conditioning experiments involving additional sensory stimuli, might contribute to a further characterization of the role of mPNs and vlPr neurons in aversive learning paradigms.

### mPNs are sufficient and required for appetitive learning

The output of mPNs is required for the retrieval of short-term sucrose-reinforced appetitive memories. By transiently blocking synaptic output from MZ699+ neurons, we show that activity of these neurons is necessary for the retrieval of 3h olfactory appetitive memory. Although the enhancer line MZ699-GAL4 labels, in addition to mPNs, 3-5 GABAergic neurons at the center of the MB calyx and a cluster of vlPr neurons (Strutz et al., 2014; Wang et al., 2014), we propose the effects seen here recapitulate those of mPNs. Previous studies have shown that vlPr activation results in avoidance behavior (Huoviala et al. 2020; Mohamed, 2019), making it unlikely that the memory deficits observed are due to the blocking of the activity of these neurons. Unfortunately, due to technical reasons, the function of GABAergic neurons in the MB calyx could not be examined. These neurons, however, do not receive evident olfactory input from the AL, making their involvement in the observed behavior unlikely.

Several studies have shown that a subset of dopaminergic (i.e. DPN-PAM neurons) and octopaminergic neurons are required for learning and retrieval of appetitive memories (Burke et al., 2012; Huetteroth et al., 2015; Liu et al., 2012; Yamagata et al., 2015). A recent study reported the presence of octopaminergic, mPNs with inputs in the AL and outputs in the LH (Bates et al., 2020). While we cannot entirely exclude that *MZ699-GAL4* labels these octopaminergic neurons, it seems unlikely that they are involved in the behavioral phenotype we report. Artificially activation of all octopaminergic neurons during odor presentation generates a memory that is short-lived: flies tested at the same time point used in our experiments (i.e. 3 hr) show no memory trace (Burke et al., 2012). Furthermore, in calcium imaging experiments we show that activity of mPNs correlates with behavior, in particular with neurons mediating odor attraction (i.e. LH-PM) that specifically show potentiation 3 hours after appetitive conditioning. Therefore, both physiological and behavioral data are consistent with an activity increase of mPNs following appetitive conditioning. This argues in favor of a model in which output from GABAergic mPNs is required for the retrieval of appetitive memories. The underlying tenet in these results is that a circuit with no apparent connection to the MB is involved in olfactory learning and memory. Such a scenario is further supported by experiments where we artificially implant memories in the fly. Interestingly, non-associative plasticity in form of olfactory habituation has also been shown to arise at the AL level mediated by inhibitory LNs (Das et al., 2011).

Artificial activation of MZ699+ neurons promotes approach-behavior: odors neutral in valence become attractive when their presentation is coupled to the activation of MZ699+ neurons. Furthermore, pairing the activation of MZ699+ neurons with a 2 min. odor presentation results in the formation of a short-term appetitive memory. Notably, the acquisition of this memory trace only occurs in starved flies, indicating that these neurons might code for the nutritional and not the sweet value of the appetitive reward (Burke and Waddell, 2011). Besides demonstrating that MZ699+ neurons are involved in the acquisition of appetitive memories, these data also indicate that GABAergic neurons are part of the reward circuitry, a role that so far has been attributed to dopaminergic and octopaminergic neurons only (Liu et al., 2012; Yamagata et al., 2015). This, however, does not exclude the possibilty that mPN activity drives pathways that end in activating dopaminergic neurons in the MBs, such as PAM-DANs (Cognigni et al., 2018).

### Interplay between LH and MB circuits

Some questions remain open. For instance, whether this parallel PN pathway is indeed MB-independentor whether MZ699+ neurons are somehow downstream or upstream of the MB circuits needs to be addressed in future experiments. If mPNs are downstream of the MB, output neurons from this brain center might modulate the activity of mPNs. Notably, manipulating the activity of mPNs has a direct impact on associative learning. Given that mPNs are also part of the first level of olfactory processing, it may be that neurons collecting information from mushroom body output neurons (MBONs) serve as top-down input to the AL (Bates et al., 2020). Indeed top-down modulation occurs in the olfactory systems of vertebrates (Adams et al., 2019; Aqrabawi et al., 2016; Chen and Padmanabhan, 2020; Kiselycznyk et al., 2006). There, centrifugal neurons, projecting from the cortex, specifically target inhibitory neurons in the olfactory bulb (Boyd et al., 2012). In *Drosophila*, several centrifugal neurons have been identified (Tanaka et al., 2012) and stimulating the MB modulates the activity of neurons in the AL (Hu et al., 2010). However, whether top-down modulation plays a role in olfactory tuning and behavior remains to be elucidated in future experiments. In our study, silencing one of these centrifugal neurons, the CSDn, did not impair aversive learning. Whether this neuron is required for appetitive conditioning or whether other centrifugal neurons are involved needs to be determined. Alternatively, it is also possible that the activity of mPNs reflects the coordinated action of the CS and US converging on them. In invertebrates, simultaneous presentation of an odor and a electric shock (or reward) leads to an increase in intracellular cAMP that depends on the activity of a protein that serves as a coincident detector (Davis, 1996; Dudai et al., 1983; Mons et al., 1999). Both the adenylyl cyclase encoded by the gene rutabaga and NMDA receptors have been proposed as such detectors (Dudai, 1988; Xia et al., 2005). Though these proteins are preferentially expressed in the mushroom bodies (Han et al., 1992), cAMP-mediated synaptic plasticity in other brain areas is poorly understood. It can well be that learning directly modulates the activity of mPNs through the action of a coincident detector.

In conclusion, we propose a model in which mPNs, not only mediate approach-behavior, but are also part of the reward circuit and in which learning modulates the innate representation of odor valence, encoded by mPNs (**Figure 8**). Olfactory learning can thus bidirectionally modulate odor-evoked responses of mPNs that, in the case of aversive conditioning, requires glutamate and serotonin signaling. These changes have opposing effects in third-order vlPr neurons, downstream of mPNs, where the decrease in GABA release from mPNs increases their activity. Finally, output from MZ699+ neurons is required for the retrieval of appetitive olfactory memories and, in the presence of odor, it is sufficient to generate new appetitive memories. Taken together, our results provide new perspectives on the neural circuits involved in the processing of learnt olfactory information and suggest that circuits underlying innate behavior are part of the engram.

**Supplementary Figure 1.**
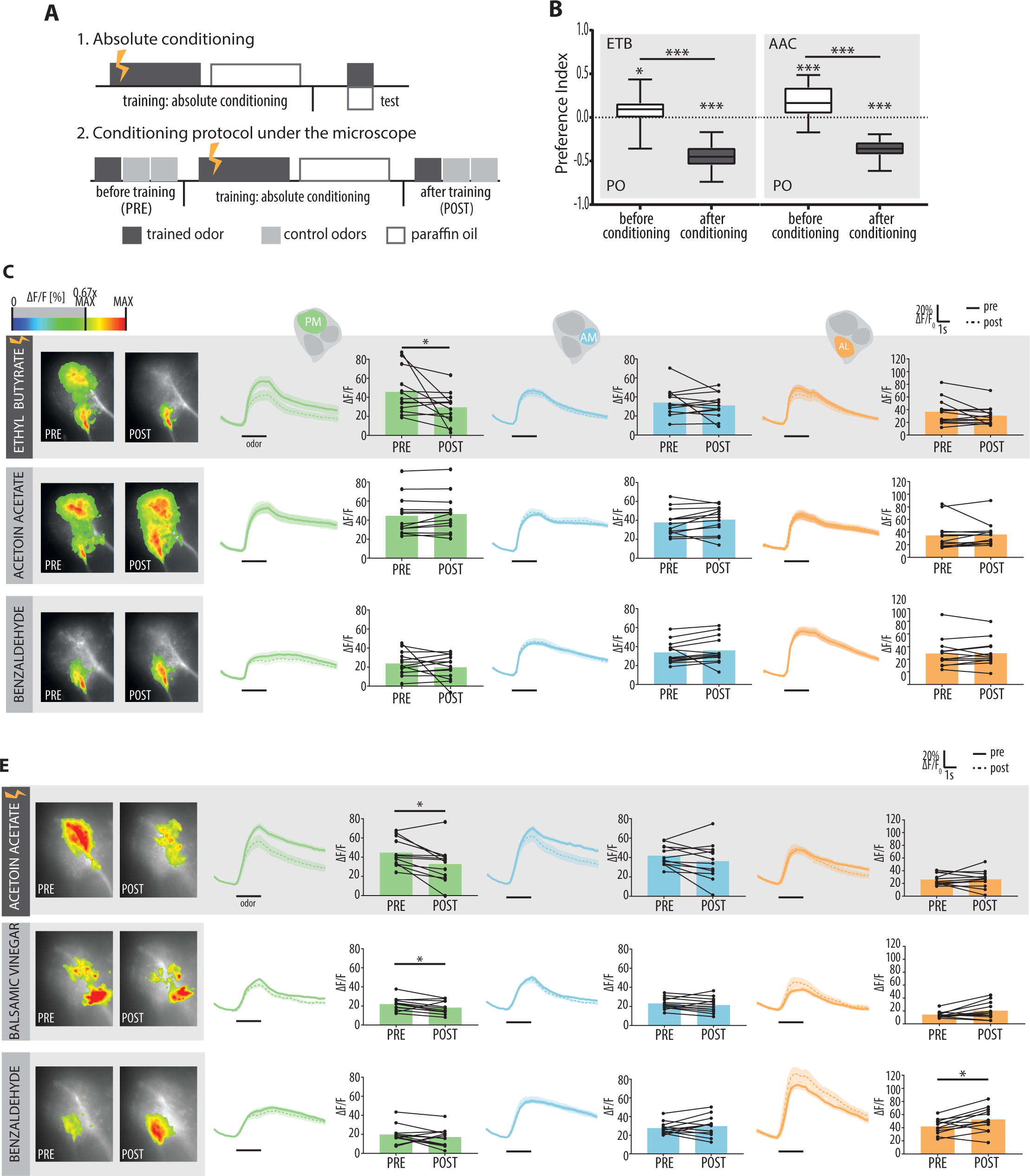
Absolute aversive conditioning with attractive odors leads to reduced odor responses in mPNs for the learned odor. (A) Training protocols for absolute olfactory associative learning used in behavioral experiments and functional imaging recordings. (B) Odor preference for the odors ethyl butyrate (ETB) and acetoin acetate (AAC) in naïve flies (before conditioning) and after absolute conditioning in a T-maze choice situation. Flies show a significant aversive response to the attractive odors after training (innate preference: n= 26 (ETB), n= 20 (AAC); after conditioning: n=20 (ETB), n= 12 (AAC); *p<0.05, ***p<0.001; genotype: *MZ699-GAL4; UAS-GCaMP6s*. Asterisks denote one sample t test against 0 or unpaired Student t test. (C and D) Functional imaging of flies expressing GCaMP6s in mPNs using *MZ699-GAL4*. Aversive conditioning leads to a significant decrease in the posterior-medial (PM) domain only for the trained odor (ethyl butyrate or acetoin acetate, respectively). Left: Representative false-color coded images showing odor-induced Ca^2+^ signals before (PRE) and after (POST) absolute conditioning for the trained and control odors. Right: time traces of Ca^2+^ signals averaged over all flies (±SEM) for each functional LH domain (PM, AM, AL) and each odor tested. Black bars represent 2 s odor stimulation. Bar plots give mean activity (ΔF/F) after odor stimulation; individual flies are given by single dots and lines (D: n=13, E: n=11, paired Student t test, *p<0.05).

**Supplementary Figure 2.**
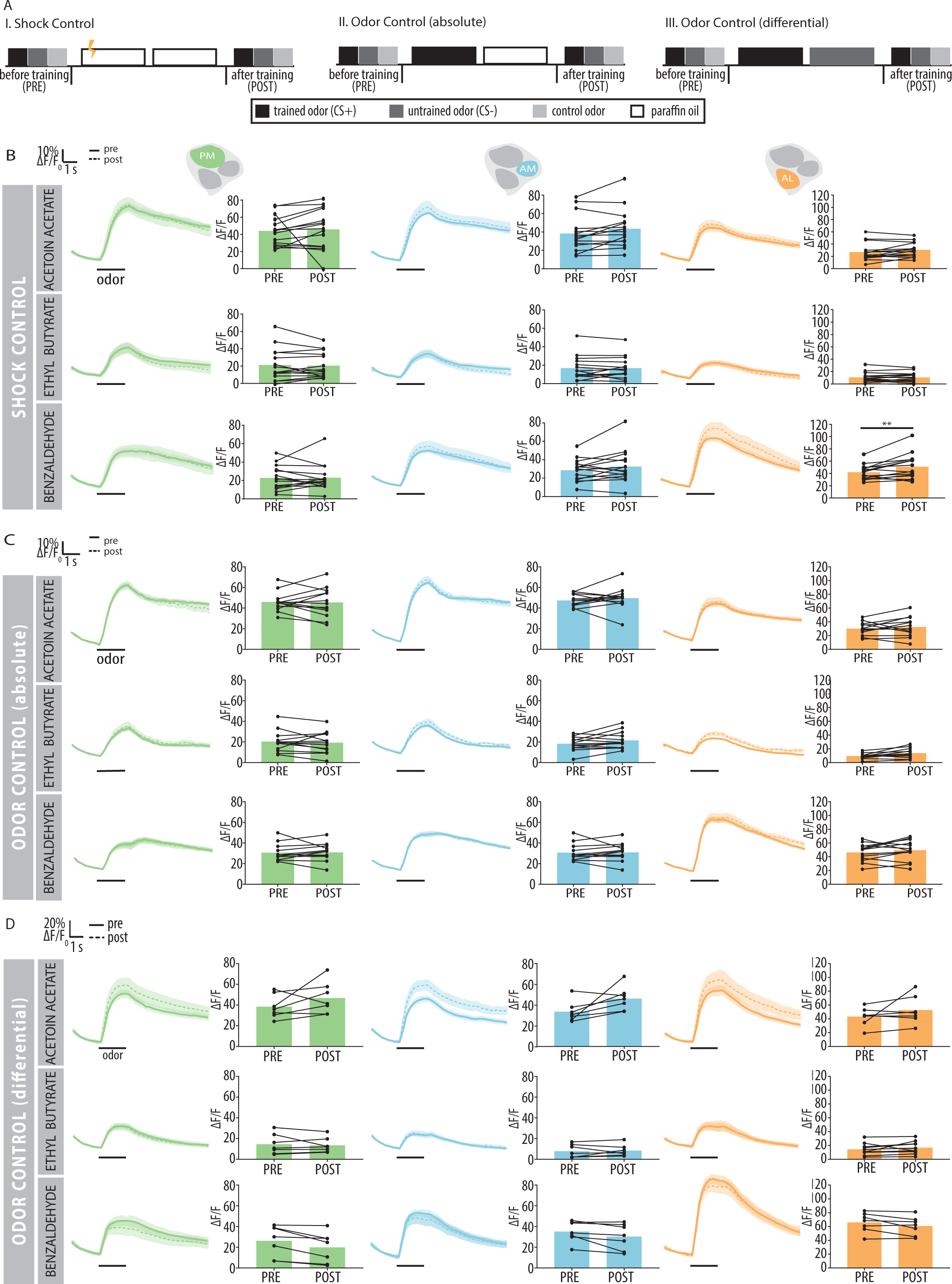
Odor-evoked Ca^2+^ changes in mPNs requires temporal overlap of CS+ and punishment. (A) Flies were conditioned using an unpaired conditioning protocol in which the electric shock (US) was delivered 2 min prior to odor (i.e. ethyl butyrate) onset. (B) Average odor-induced Ca2+ activity in each functional domain (PM, AM, AL) for each odor used (n=12, Student paired t test, ns).

**Supplementary Figure 3.**
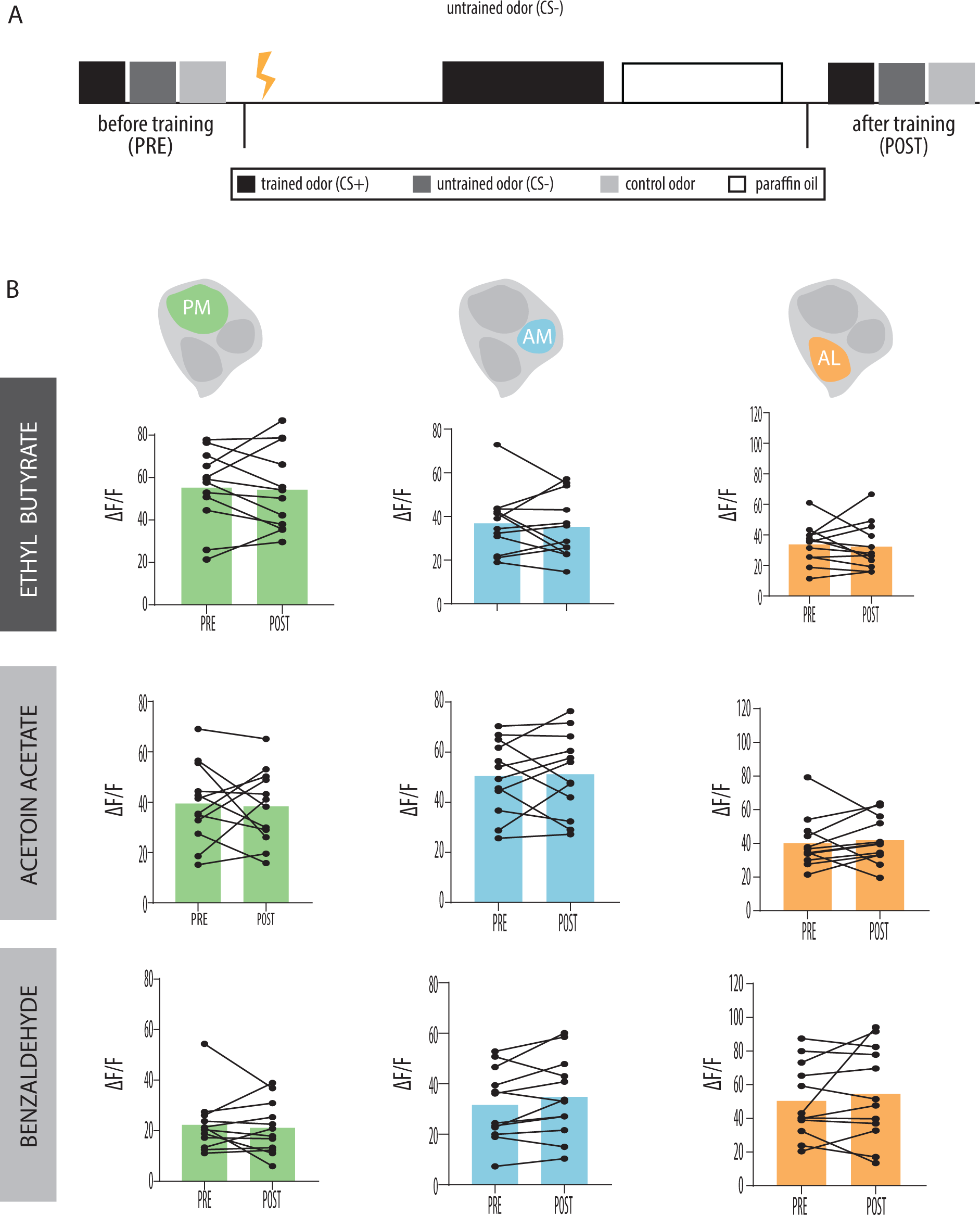
Odor presentation and shock application alone do not result in odor-evoked Ca^2+^ changes in mPNs. (A) Schemes illustrating the mock training protocols in which (I) an electric shock was applied in absence of odor (shock control) or (II, III) the odor was presented without negative reinforcement (odor control). (B, C, D) Time courses of odor-evoked GCaMP responses in each functional domain (PM, AM, AL), before (pre) and after (post) treatment. Traces represent mean odor response (solid line) and standard error of the mean (shading). Bars show average peak response upon odor presentation (n=13, paired student t test, **p<0.01).

**Supplementary Figure 4.**
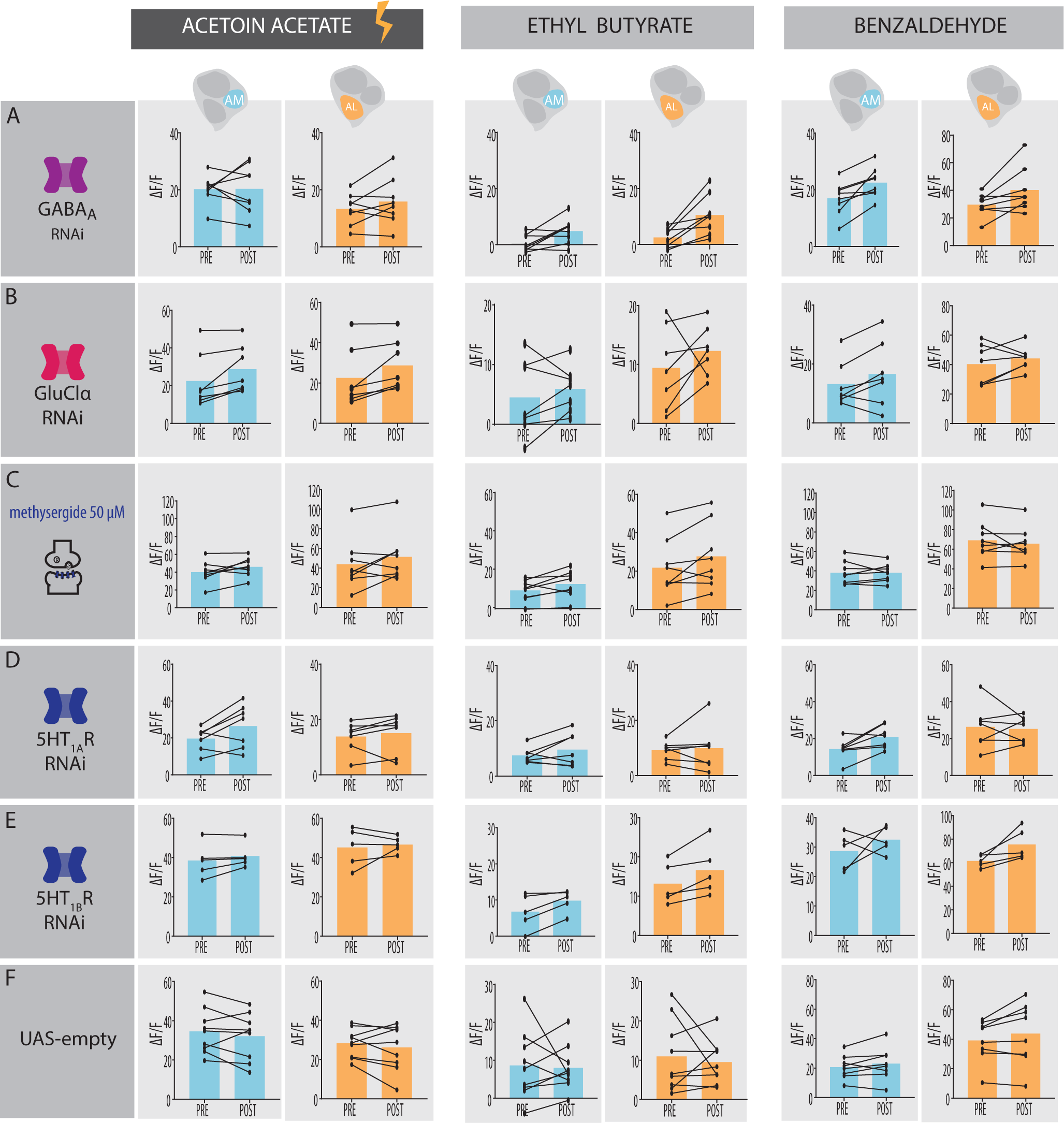
Learning-induced response decrease in mPNs after training requires glutamate and serotonin receptor activity. Odor-evoked peak responses in mPNs AM and AL domains before (PRE) and after (POST) aversive differential conditioning, from flies presenting the genetic or pharmacological manipulation indicated on the left: (A) Expression of *UAS-RdliG-GABA_A_ RNAi* was used to downregulate GABA-A receptor expression in MZ699+ neurons (n=8, ns). (B) Downregulation of glutamate receptors in MZ699+ neurons via expression of *UAS-GluClα RNAi* (n=7, ns). (C) Pharmacological inactivation of serotonin (5HT) receptors using the receptor antagonist methysergide (n=8, ns, ns). (D, E) Downregulation of inhibitory 5HT receptor subunits using either *UAS-5HT1A R*NAi or *UAS-5HT1B RNAi* in MZ699+ neurons (5HT1A: n=5; 5HT1B: n=7, ns). (F) Flies expressing the control transgenic construct *UAS-empty RNAi* (n=8, ns). All statistics are paired Student t test (also see Figure 5).

**Supplementary Figure 5.**
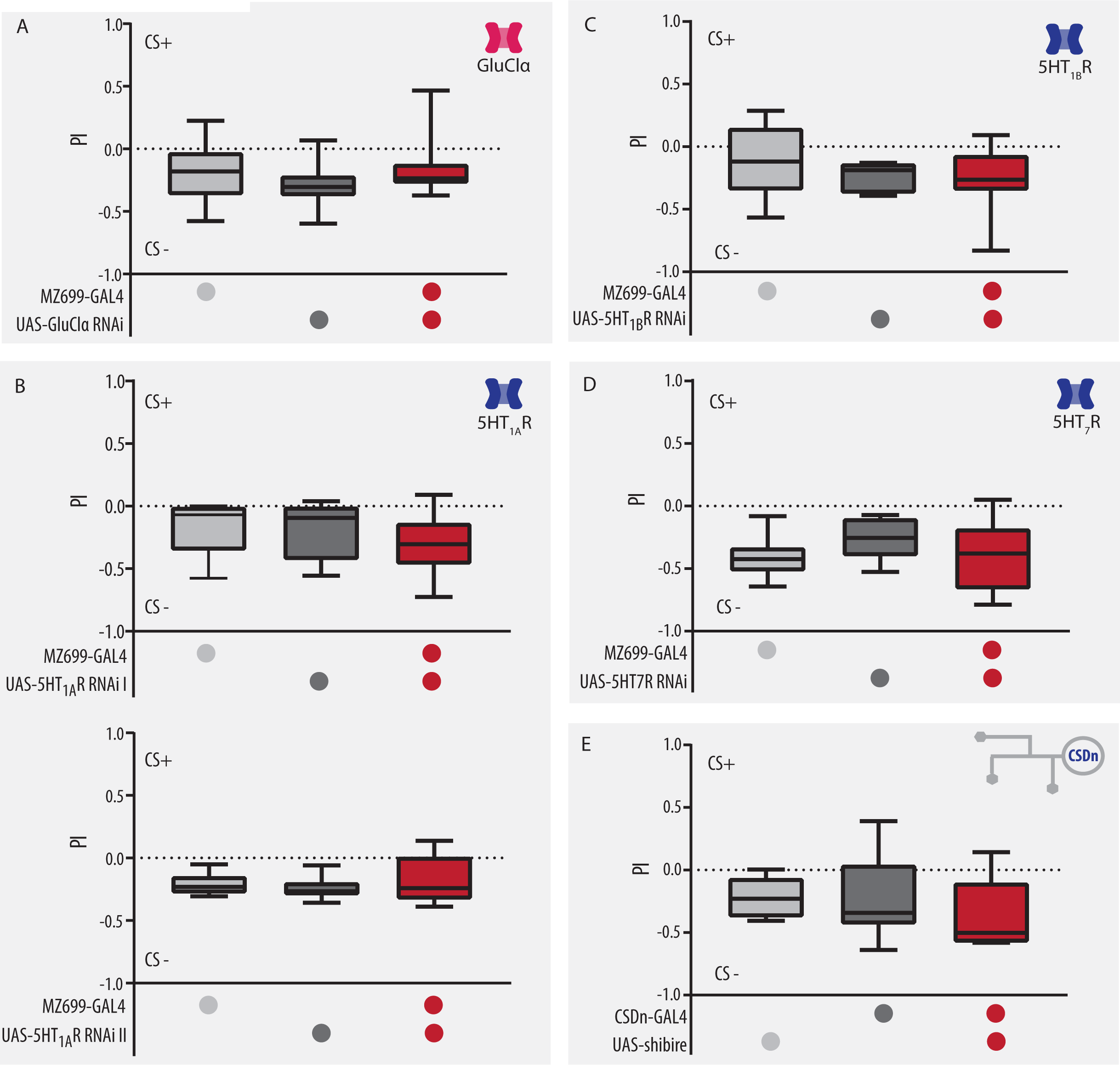
Downregulation of glutamate and serotonin receptor subunits in MZ699+ neurons does not impair aversive differential associative learning. Flies do not show deficits, respective to their genetic controls, in short term memory (STM) (as shown by the performance index (PI) after training) when MZ699+ neurons express RNAi against the receptors (A) GluClα (n=16-17, ns), (B) 5HT_1A_, using line I (n=5-10, ns) or line II (n= 7-10, ns) (C) 5HT_1B_ (n= 8-12, ns), and (D) 5HT7R (n= 6-8, ns). In (B), line I expresses the same RNAi transgene used in calcium imaging experiments (see Fig 4). (E) Blocking synaptic output from the serotonergic CSD neuron using thermo-sensitive *shibire^ts1^* during training and retrieval does not impair STM (n= 5-7, ns). All statistics are one-way ANOVA followed by a Tukey’s multiple comparisons post-hoc test.

**Supplementary Figure 6.**
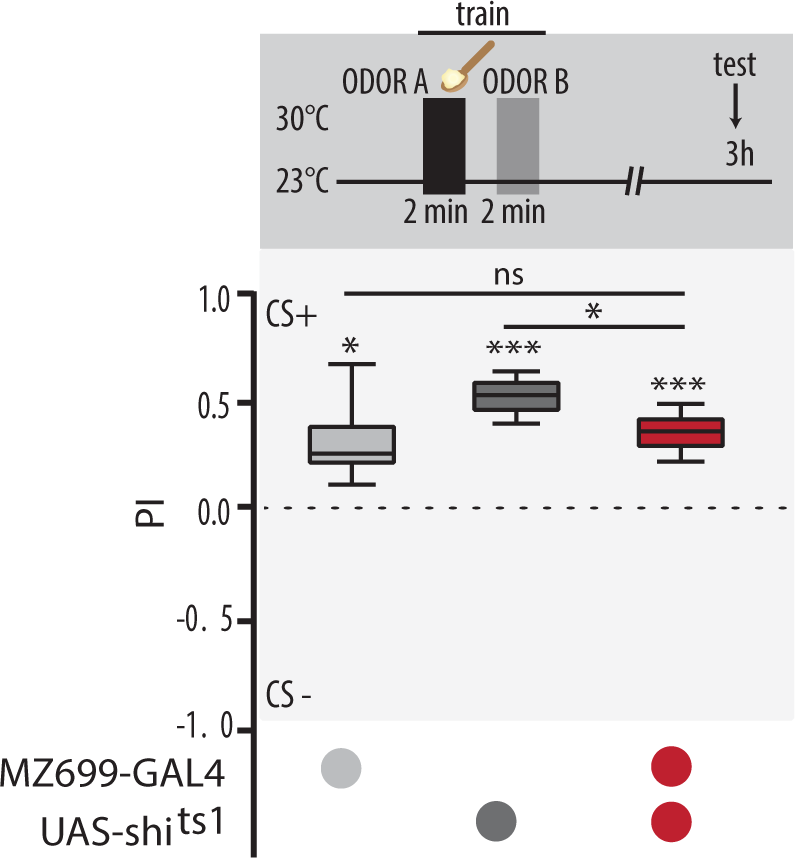
MZ699+ neurons are required for the retrieval of appetitive memories and are sufficient to generate an appetitive memory. (A) Top, scheme illustrating training protocol for differential appetitive olfactory conditioning with 3-octanol or 4-methylcyclohexanol as CS+. Flies were trained and tested 3 h later at the permissive temperature. Bottom: Flies expressing shibire^ts1^ show a partial reduction in memory retrieval, but are still significantly attracted to the trained odor (n= 6-8, one-way ANOVA with Tukey’s multiple comparisons post-hoc test; one sample t test against 0; *p<0.05, ***p*<*0.001).

